# A review of the International Seabed Authority database DeepData: challenges and opportunities in the UN Ocean Decade

**DOI:** 10.1101/2022.10.14.512288

**Authors:** M. Rabone, T. Horton, D. O. B. Jones, E. Simon-Lledó, A. G. Glover

## Abstract

There is an urgent need for quality biodiversity data in the context of rapid environmental change. Nowhere is this more urgent than in the deep ocean, with the possibility of seabed mining moving from exploration to exploitation, but where vast knowledge gaps persist. Regions of the seabed beyond national jurisdiction, managed by the International Seabed Authority (ISA) are undergoing intensive mining exploration, including the Clarion-Clipperton Zone. In 2019 the ISA launched its database ‘DeepData’, publishing environmental (including biological) data; and since June 2021, DeepData records have been harvested by OBIS (Ocean Biodiversity Information System) via the ISA node. Here we explore how DeepData could support biological research and environmental policy development in the CCZ (and wider ocean regions); and whether data are Findable, Accessible, Interoperable and Reusable (FAIR). Given the direct connection of DeepData with the regulator of a rapidly developing potential industry, this review is particularly timely. We found evidence of extensive duplication of datasets; an absence of unique record identifiers and significant taxonomic data quality issues, compromising FAIRness of the data. The publication of DeepData records on the OBIS ISA node has led to large-scale improvements in data quality and availability. However, limitations in usage of identifiers and issues with taxonomic information were also evident in datasets published on the node, stemming from mis-mapping of data from the ISA environmental data template to the data standard Darwin Core prior to data harvesting by OBIS. While notable data quality issues remain, these changes signal a rapid evolution for the database and significant movement towards integrating with global systems through usage of data standards and publication on global aggregators. This is exactly what has been needed for biological datasets held by the ISA. We provide recommendations for future development of the database to support this evolution towards FAIR.

## 2. Introduction

The need for high quality biodiversity data is abundantly clear in the face of the biodiversity crisis, with numerous pressures impacting species, including climate change (1). Such data are essential for understanding ecosystems, detecting and monitoring anthropogenic impacts and developing effective environmental policy. To be usable for both research and policy, it is important that data meet criteria of being FAIR, or Findable, Accessible, Interoperable and Reusable (2). For example, FAIR biodiversity information can be fed into frameworks for monitoring and observation, such as Essential Ocean Variables (EOVs) and Essential Biodiversity Variables (EBVs); and utilised in environmental policy (3, 4). However, major gaps in coverage of global biodiversity data across thematic and geographical areas have been identified (5, 6). Further, the biodiversity data landscape is highly heterogenous, with varying degrees of data integration and exchange (7, 8, 9, 10). This landscape is characterised by a multitude of databases, some highly specialised, by theme, region, taxon or similar (e.g. Fishbase; www.fishbase.org), and some broad, global aggregators, e.g. GBIF, the Global Biodiversity Information Facility (https://www.gbif.org).

Relevant data types in biodiversity include taxonomy, occurrence, environmental, and genetic/genomic data (8; Figure 1). Biodiversity databases often specialise by data type, e.g. the World Register of Marine Species (WoRMS^1^) in taxonomy, as a checklist and classification of marine taxa (11, 12, 13); and exchange information, e.g. the ocean data aggregator OBIS (Ocean Biodiversity Information System) which specialises in occurrence and environmental data, and utilises the WoRMS taxonomic backbone (14, 15). Global data standards such as Darwin Core (DwC) administered by Biodiversity Information Standards (TDWG) allow for data interoperability and exchange (16). In addition to data standards such as DwC, there are many relevant standardisation efforts. For example, the Ocean Best

**Figure 1.**
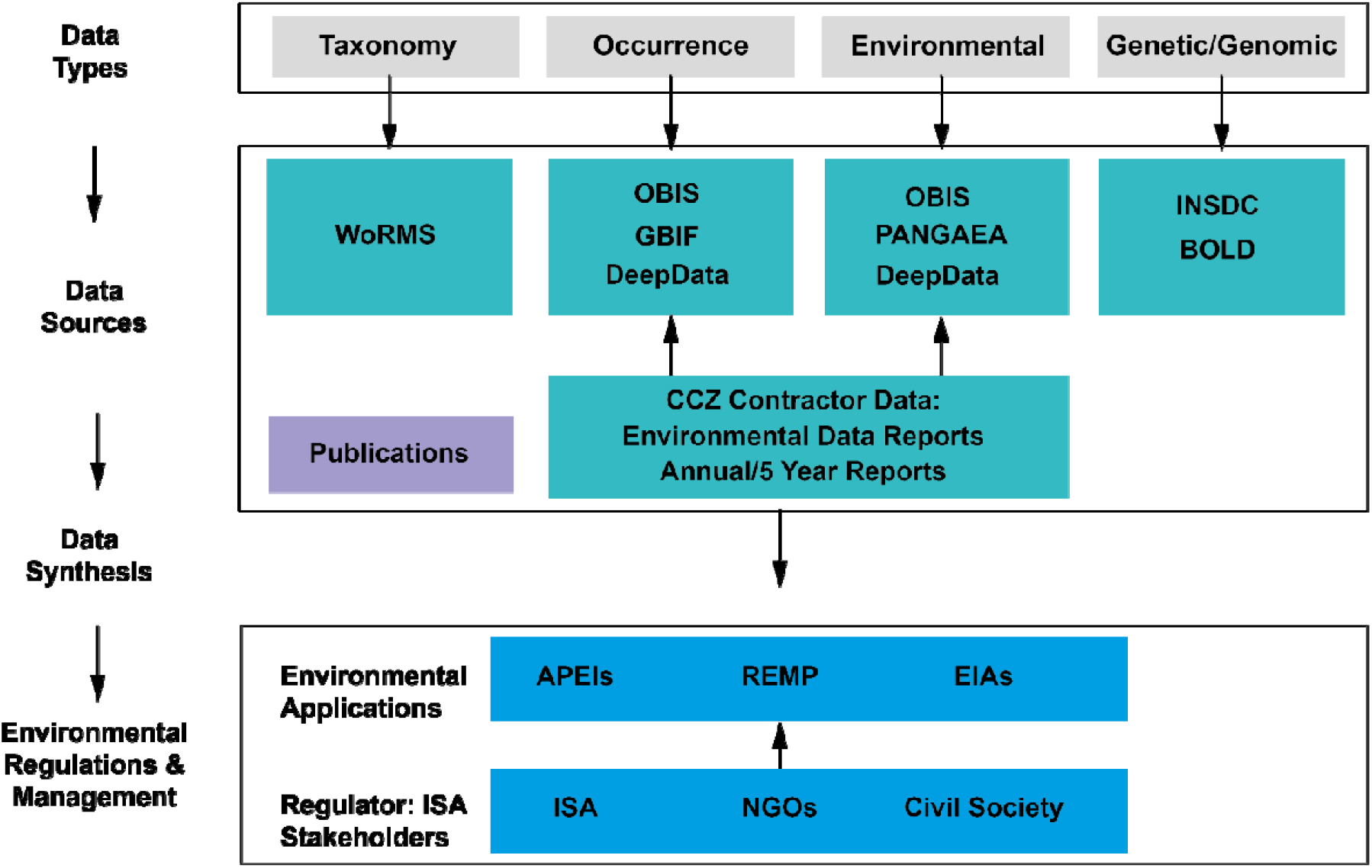
The Clarion Clipperton Zone biodiversity data landscape, showing relevant key data types: taxonomy, occurrence, environmental and genetic/genomic data; key data sources, databases, publications, and contractor data; and how these data, once synthesised in publications and meta-analyses and could contribute to environmental management applications, with input by the regulator, the ISA (International Seabed Authority) and wider stakeholders. Key databases as listed include the following: WoRMS: World Register of Marine Species; OBIS: Ocean Biodiversity Information System; PANGAEA: Data Publisher for Earth & Environmental Science; INSDC: International Nucleotide Sequence Database Collaboration; BOLD: Barcode Of Life Data System. Environmental applications: APEIs: Areas of Particular Environmental Interest; REMP: the Regional Environmental Management Plan; and EIAs: Environmental Impact Assessments. Thankyou to the Nautilus biodiversity data working group where a Miro board sketch inspired elements of the figure.

Practices System (OBPS) under the auspices of the Intergovernmental Oceanographic Commission (IOC) provides a platform for best practices with a ‘semantic’ approach, linking relevant protocols (17, 18, 19). However, adoption of standards and best practice is variable (20, 8). Key challenges include treatment of taxonomic information, a long-standing issue in biology (21, 11, 12); and problems with validity of identifiers, compromising data exchange, traceability, and contributing to duplication (7, 22, 20, 23, 24, 25).

Nowhere are these challenges more apparent than for deep-sea biodiversity data, where most species are undescribed (26, 27, 11). Extensive usage of morphospecies names, temporary names given to species prior to formal description (for example, 28; 29, 30), or ‘open nomenclature’ *sensu* Horton et al., (11) and Sigovini et al., (31) compound the existing challenges in taxonomy. The open ocean and the deep sea represent key information gaps in global biodiversity data coverage (5, 32, 33). However, regions of the deep ocean are undergoing intensive exploration for mining of polymetallic nodules, in particular the Clarion-Clipperton Zone in the Central Pacific. The deep seabed is managed by the International Seabed Authority (ISA), regulator of mineral-related activities; a body established under UNCLOS, the United Nations Convention on the Law of the Sea. As part of the mineral exploration process, the ISA requires the holders of exploration contracts to collect and make available biological data to improve the understanding of deep-sea ecosystems and the impacts of potential deep-sea mining activities (34).

The need for a central ISA database was formally identified by the Legal and Technical Commission (LTC) of the ISA in 2002 (ISBA/8/C/6). Several LTC recommendations were made during the period of 2002-2019^2^, and DeepData was developed as a further iteration of the previous Central Data Repository, which had been primarily focussed on mineral resources. In 2019, the ISA launched the public database DeepData as a repository of deep-seabed related data collected by contractors and related parties (e.g. research organisations conducting surveys) in the Area (https://data.isa.org.jm/isa/map). The database holds both geological data, categorised as confidential, and publicly available environmental data, an umbrella term for environmental and biological data in ISA parlance. DeepData is unusual in the respect that there is a direct connection of the key database, DeepData with the regulator, ISA, and that the main data providers to the database are contractors undertaking exploration of mineral resources in the Area, albeit working directly with the scientific community. The microcosm of the CCZ data landscape can however illustrate processes of how biodiversity data types are collated and subsequently published from a range of sources (Figure 1). Further, given the direct connection of the regulator and database, it also illustrates how these data could be synthesised and applied to environmental management, for example in developing tools such as the Regional Environmental Management Plan (REMP) or the design of Areas of Particular Environmental Interest (APEIs; see Figure 1 and Smith et al., (35)).

In this study, we provide the first review of DeepData, focussed on the biological data available for the most active area of seabed mining (CCZ) and include recommendations for the development of this database into the future. This work is particularly timely given DeepData has now been operational for four years; associated records are being actively pushed onto global data aggregators such as OBIS, GBIF and INSDC (International Nucleotide Sequence Collaboration) and OBIS is also now publishing DeepData records via the OBIS ISA node (3). A more critical point however is the context of the rapid recent development of deep seabed mining regulations and the urgent need to address deep-sea biodiversity data gaps both for the CCZ and other regions (36, 37). Here we conduct an assessment of the database and wider related ISA biological/environmental data management as part of a broader study where we synthesise the biodiversity and biogeographic data available from DeepData and associated databases for the CCZ (Rabone et al., in prep). The primary purpose of this review therefore is to assess the FAIRness of published biological data in DeepData, and the potential utility of the database to support both research and decision-making for environmental policy.

## 3. Materials and Methods

### Overview of DeepData and description of the online data portal

The ISA DeepData website or online data portal provides biological, geochemical, and physical data collated from expeditions arranged by contractors for the CCZ and other exploration regions. The map-based interface includes boundary data (e.g., shapefiles) depicting APEIs, mining exploration contract areas, reserved mining exploration areas, and research sample data. The datasets currently held in the database include: biological and geochemical analyses from samples collected using box corers, epibenthic sledges, multiple corers, ROVs and benthic trawls; navigational information from expeditions; current meter recordings; and water column and water sampling data. The DeepData web interface has two windows, ‘HOME’ with a map view, and ‘MAP OPTIONS’, with 6 tabs: ‘Layers, ‘Search’ ‘CTD’, ‘Photo/Video Gallery’, ‘Library’, and ‘Docs’ on the left-hand side, with the map on the right (S Figure 1; https://data.isa.org.jm/isa/map/). Options to select biological data by category, on the ‘MAP OPTIONS’ window, ‘Layers’ tab are as follows: ‘Contractors - Mineral Type’^4^; ‘Contract Status’ (all/active/extended), ‘Sponsoring State’; ‘Mineral Type’ (Cobalt Rich Ferromanganese Crust (CRFC)/Polymetallic Nodules (PMN)/Polymetallic Sulphides (PMS)); and ‘Location’ (Central Indian Ocean/Central Indian Ridge and Southeast Indian Ridge/ Clarion-Clipperton Fracture Zone/Indian Ocean/Indian Ocean Ridge/Mid-Atlantic Ridge/Rio Grande Rise/South Atlantic Ocean/Southwest Indian Ridge/Variable - PMN Reserved Areas/Western Pacific Ocean). Options to search and download data are on the adjacent ‘Search’ tab, and under ‘filter by data type’ is a dropdown menu to select first, data type: ‘Biological’, or ‘Environmental Chemistry’, and second, sampling method: ‘Point’, or ‘Trawl Line’ (S Figure 1). Here ‘Points’ equate to deployments (sampling events) collected from a particular point in space and time e.g. a box core; and ‘Trawl lines’ being those collected from sampling between two points, e.g. and ROV or via towed gear such as a Brenke Epibenthic Sledge trawl sample.

### Data Collection

Biological data were downloaded from the DeepData database web portal on the 12^th^ of July, 2021. The data selection was conducted as follows: ‘Layers’ tab: ‘Mineral Type’: ‘Polymetallic Nodules’, ‘Location’: ‘Clarion Clipperton Fracture Zone’, Search tab, ‘Biological data’, ‘Point’, and to export the data, ‘export query’ (S File 1A). The same search procedure was run again with the ‘Biological Data’ option as ‘Trawl line’ for the trawl-collected data (S File 1 B). For contextual spatial data, all mining exploration contract areas, both active and reserved, and APEI shapefiles were downloaded from the ISA database (https://www.isa.org.jm/minerals/maps) and combined into one shapefile in QGIS. Coordinates for a polygon covering the entire CCZ including the combined shapefile were established: (in decimal degrees, longitude/latitude): northwest -164.01462, 15.70629; southwest -155.04998 -5.51238; southeast -101.9181 6.05623; northeast -117.66088 23.72549. DeepData records have been harvested by OBIS since June 2021 (https://obis.org/node/9d2d95be-32eb-4d81-8911-32cb8bc641c8). OBIS occurrence data were downloaded as a Darwin Core file on the 12^th^ of July, 2021 using the ‘occurrence’ function in the robis package (Provoost & Bosch, 2017), with the CCZ polygon as delineated above, for all depths.

### Data Processing and Analysis

#### Data restructuring and general data processing

Data were processed and analysed in R, version 4.0.2 (2020-06-22) “Taking Off Again” (R Core Team, 2020). General quantitative and qualitative observations as well as structured notes were made for analysis. Preliminary investigations of the database export showed that the records (or observations) were distributed both across columns and rows; rather than one record per row (38). The data were restructured to one record per row using the ‘spread’ function in R from the tidyverse package (39). The separate ‘Point’ and ‘Trawl Line’ data downloads were combined into the same dataset (S File 1C). As the data fields varied between the two datasets, e.g. ‘actual latitude’ in the ‘Point’ data, and ‘startLatitude’ and ‘endLatitude’ in the ‘Trawl Line’ data, fields were harmonised. For coordinates and depth, the end-point was used, i.e. ‘endLatitude’ was mapped to ‘actualLatitude’, to allow the datasets to be combined (S File 1C, D). Initial assessments of the data also found that the database output did not contain a record identifier, or a unique key in any format, primary, composite or other. To examine the data, a composite key was created, combining the DeepData identifier fields for contractor, station and specimen (‘ContractorID’ + ‘StationID’ + ‘SampleID’). It was checked for duplicates, and none were found. Data columns were checked and edited where necessary (e.g. for depth, missing values were listed as -9, these were replaced with ‘NA’). Where possible this was scripted in R, where multiple entries for character variables were present, this was done in Microsoft Excel 365 on a copy of the data column, renamed with the suffix “_ed” (S File C, D).

#### Geographic Mapping

Contractor sub-areas were mapped in QGIS and data revised to reflect actual geographic areas, rather than origin of records, i.e. ContractorID (name of contractor submitting data) as these were not equivalent. All OBIS records were mapped together with the CCZ shapefile, using the following R packages: GADMTools, sp, spData, spatialEco, maptools, rgdal and rgeos. The records were then sub-selected by depth, with depths of 3000m and greater included. Some records without depth values were present, those falling within or near the CCZ shapefile were reviewed and included if valid, for example if a benthic species/taxa associated with a publication and a benthic collection method e.g. a box core sample; and/or a relevant reference in ‘datasetName’ or ‘associatedReferences’ column. The DeepData records published on OBIS were sub-selected from general OBIS records (distinguished as recorded as owned by the ISA in the Darwin Core ‘accessRights’ field; S File 2).

#### Taxonomic data

Initial examination of taxonomic information found extensive inconsistent recording of names, e.g. misspellings, mis-formatting (e.g. escaped newlines) and mis-recording, e.g. class names recorded in the family field. This is typical in many new species occurrence databases that are not linked to a taxonomic source. No DwC equivalent field to ‘scientificName’ was present, i.e. the lowest taxonomic level identification of the specimen referenced in a given record. To allow data to be analysed for the parallel study (Rabone et al., in prep), this field was added, populated with the lowest taxonomic level identification present per record. If a name was noted with question mark, recorded with the qualifier incertae sedis or written as two names, the next highest taxonomic level recorded was added as the scientific name. For example, if two Family names were present, indicating a level of uncertainty in the identification, or an identification qualifier such as incertae sedis was recorded in notes, then the Order was recorded as the scientific name. Preliminary investigations showed significant numbers of morphospecies names, and/or ‘open nomenclature’ designations, e.g. names recorded with qualifiers, such as cf. (11, 31). Where open nomenclature designations were provided (in the DeepData field ‘putative species name or number’), a scientificName was also recorded, mapped to the lowest taxonomic level identification above species level. If a species name (i.e. specific epithet) was present in the ‘putative species name or number’ field, then the genus name only was recorded in the scientificName field. The taxonomic information was cleaned using ‘taxonMatch’ in WoRMS, a QA/QC function on the website where scientific names can be validated against the database (www.marinespecies.org). Resulting names were cross-referenced, any usage of unaccepted names recorded, and corresponding accepted names added to the newly created ‘scientificName’ field. If no match was found on WoRMS, the original name was retained. Any qualifiers recorded with a name, e.g. ‘cf.’ In the genus field were mapped to a separate identification qualifier field and the taxonomic level of the qualifier recorded. A sample of contractor data submissions was requested from the ISA for insight into both ISA data mapping and processing and contractor data recording. A selection of records from six contractors from annual data reporting submissions from 2015-2017 were provided, and datasets were harmonised and processed into one file (S File 3). Structured notes were made on taxonomy fields both for the published records and the unprocessed contractor data files, e.g. on spelling errors, formatting issues and similar, for general context and comparison.

## 4. Results

### 4.1. Data structure of database output

The data export from DeepData of biological ‘Point’ data from the 12^th^ of July consisted of a dataset of dimensions: 98,1483 rows, 48 columns. Post data restructuring to one observation per row resulted in a file of 52,177 rows, 56 columns. The data export of ‘Trawl Line’ data consisted of a much smaller dataset of 941 rows and 49 columns, restructured to 45 rows. The two files were then combined to produce a final dataset of 52,222 rows, 56 columns (S File 1C). As the wider study was examining benthic metazoa only, records of non-metazoans, such as xenophyophores, or records without taxonomic information were removed. This resulted in a final dataset for analysis encompassing 40,518 rows, 56 columns (S File 1D, also used in the parallel study, i.e. Rabone et al., in prep). The distinction between ‘Points’ and ‘Trawl Line’ for records in DeepData was incomplete, with numerous trawl-collected records evident across multiple datasets e.g. benthic plankton trawls and epibenthic sledge-collected samples which would in theory both be categorised as ‘Trawl Line’, present in the ‘Point’ dataset (S File 1A-D). The ‘Trawl Line’ data in the database output contained a sole dataset of 45 records from a single dataset, but >8000 records in total in DeepData would fall into a ‘Trawl Line’ classification (e.g. collected by an epibenthic sledge, benthic trawl, AUV or ROV). This distinction is therefore unnecessary (as sampling method is recorded in a separate column), requires additional data processing, and inaccurate, as ‘Point’ data appears to be used as the default category, regardless of the actual sampling method information present.

The structure of the DeepData output had observations distributed both over rows and columns, or in both ‘wide’ and ‘long’ format (38), resulting in a similar outcome of additional data processing steps. Wide format is one record or observation per row; and ‘long’ format’ where one record or observation is split across multiple rows. All data were wide format, until the fields ‘Analysis’ and ‘Result’, where these data fields were ‘paired’, i.e. ‘Result’ data values pertain to the adjacent field ‘Analysis’, and these data were therefore structured in long format. The field ‘Analysis’ is a list of column headings, e.g. ‘Taxonomist’, ‘Taxonomist E-mail’. These headings originate from the environmental data template (S File 4A, B), and are grouped by ‘category’ field two columns to the left (e.g. for ‘Category’: ‘Taxonomist information’, column headings as recorded in ‘Analysis’ include: ‘Taxonomist’, ‘Taxonomist E-mail’ etc). The ‘Result’ field records the related data for the adjacent ‘Analysis’ field, e.g. ‘Taxonomist’ in ‘Analysis’ column, and ‘Not Reported’ in ‘Result’. The Analysis and Result columns are therefore paired, while the remainder of the table is ‘wide’ format. This is illustrated with a subset of data in S Table 1. This structure, with observations distributed both across rows and columns has produced significant redundancy in the data, only 5 columns are shown (S Table 1), but there were 48 columns in total for ‘Point’ data (and 49 for ‘Trawl Line’ data), the majority containing this redundant repeated data-39,066,594 cells in total. This redundancy will therefore multiply as more datasets added to the database. This is likely to significantly impact processing speeds. Another export option was available, ‘export pivot query’, this option has all data in wide format, but was not used in analysis as during initial exploratory investigations it appeared to differ in visual formatting only and export query was appeared to be the default format.

### 4.2. Data quality in database output

#### Taxonomy

As the database output lacked a field equivalent to the DwC term scientificName, i.e. the lowest taxonomic identification of a given occurrence record, interpretation of the identification from the available taxonomic data fields and mapping of this information to a newly created field was required. The output did not include a separate field for identification qualifier, with this information only recorded in a notes field or the actual taxonomy field/column (e.g. ‘cf. Munnopsidae’). Extensive usage of unaccepted names, misspellings and notes in taxonomic data fields was evident. This is clearly illustrated with the Phylum field, which contained 74 different entries while only 31 metazoan phyla are currently recognised. The lower the taxonomic rank however, the more variable were the data entries present. Examination and cross-referencing of unprocessed contractor files revealed that taxonomic information for all fields was published verbatim (or close-to) from contractor data submissions (S File 3), within minimal data processing evident. Where data processing of taxonomy has occurred however, it appears to have caused additional complexities, such as taxonomic designations even being changed in some cases. As an illustration, records of the annelid *Monticellina* Laubier, 1961 were present in DeepData incorrectly as *Monticellina* Westblad, 1953 (Platyhelminthes) rather than *Monticellina* Laubier, 1961 accepted as *Kirkegaardia* Blake, 2016 (Annelida). In the contractor data submissions, it was evident the relevant record was *Kirkegaardia* by comparison with the higher taxonomy columns, but the genus name was recorded as the unaccepted homonym *Monticellina* in the record. A taxon match for the genus *Monticellina* in WoRMS returns an ‘ambiguous match’ (a standard result for homonyms, pre-occupied names and similar) with the two options (*Monticellina* Westblad, 1953 and *Kirkegaardia* Blake, 2016). The DeepData name matching appears to have been carried out with reference to the lowest taxonomic level only, as the record was taxon matched to *Monticellina* Westblad, 1953 (i.e. the platyhelminth genus) rather than correctly to the annelid genus *Kirkegaardia* Blake, 2016, an error that would have been picked up if higher taxonomic ranks were cross-referenced.

#### General data quality and missing information

Several other fields also required cleaning and harmonising of data where data would match a standard set of terms, i.e. a controlled vocabulary. For example, the DeepData field ‘SampleCollectionMethod’ had variable entries, including misspellings (e.g. multi core, MUC, Multi Corer, Multi-corer). Contractors have recorded these data in variable ways in the data templates (S File 3), and like the taxonomic data, the entries had not been harmonised prior to publication. For some fields, the origin of the information present was not clear as it does not appear in the contractor templates (S File 3). For example, in the field ‘HabitatType’, approximately half the DeepData records had habitat recorded as ‘water column’, but none of the corresponding Contractor files had ‘water column’ recorded in the habitat field, or elsewhere (see S File 3). In addition, 90% of data overall were missing or incomplete for multiple fields, including key information, such as sampling method, which is critical information for analysis. For the deep-sea, size class is regarded as key information with faunal groupings generally distinguished by size (i.e. micro, meio, macro and megafauna). Data on size class (‘nominalSizeCategory’), was often missing also, despite being a required field in the data template. In some cases, omission of information has produced inaccuracies. For example, the field ‘Identification Method’ for recording how taxa were identified, text entries were present as ‘Morphological’ or ‘DNA’, but not as a combined entry, i.e. ‘Morphological and DNA’. This data recording is an artefact of an earlier iteration of the data template, where only one method could be recorded in the field and can give the impression that an identification was made with only one method even when this was not the case. As a wider point, data from the majority of cruises are yet to be published on the database, as 103 cruises have been carried out in the CCZ (ISA Secretariat, pers. comm.), but records from 24 cruises, and ten contractors in total have been published to date (Table 1; Rabone & Glover, in review). It is unclear is this is entirely due to a data backlog or if there are cases of active contractors who have not submitted data. While substantial data processing (and in some cases, interpretation) was required for taxonomy and to a lesser extent, sampling information, site data in contrast required minimal processing. Some anomalies were still evident, for example in the contractor sub-area field, a number of cases were designated as ‘OA’ (outside area) but were within the claim of that contractor (S File 1C, D).

**Table 1.**
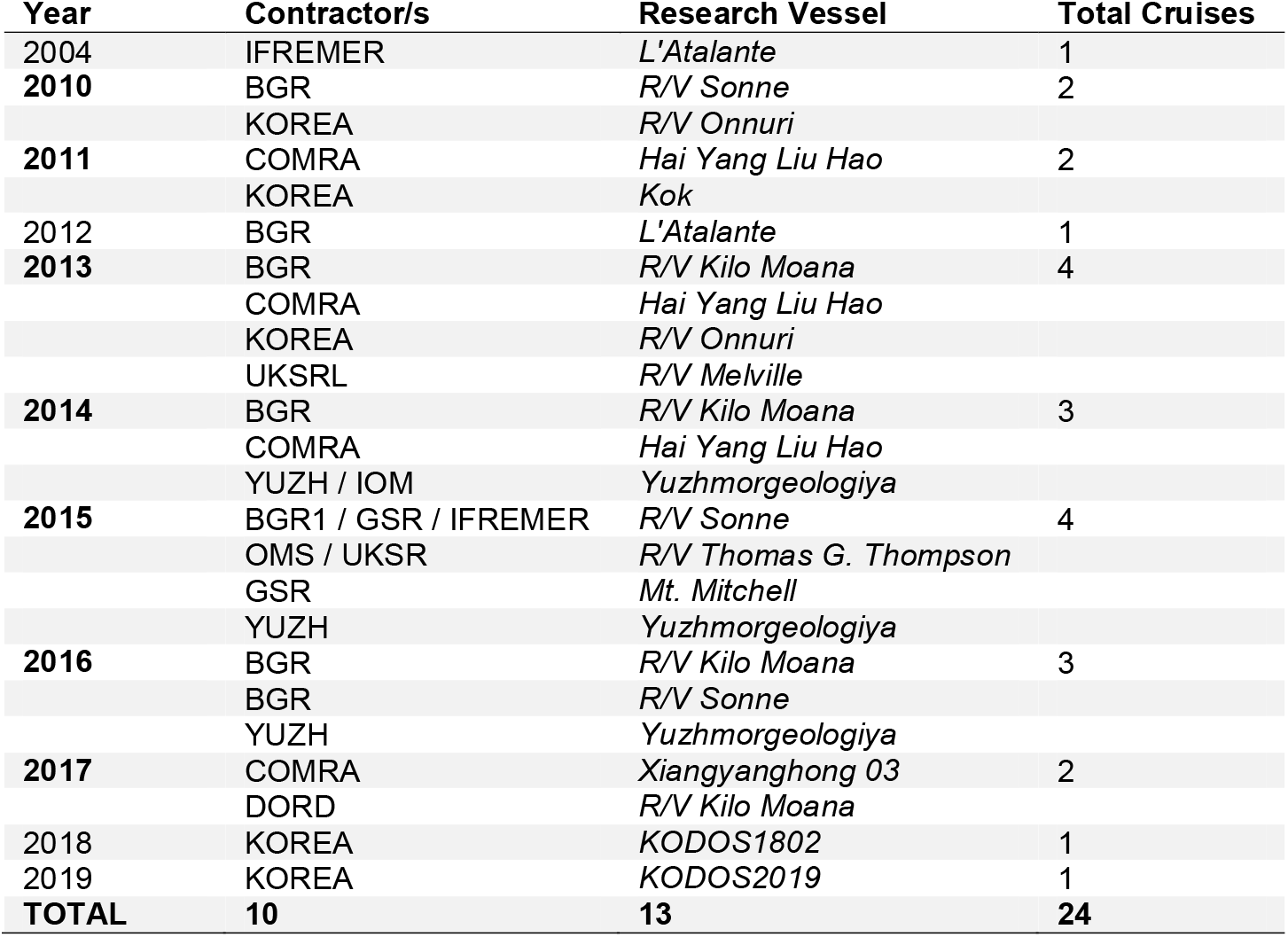
Cruises by year, contractor and research vessel in records in DeepData, as published at the time of the study (12^th^ of July, 2021). Years in bold-where datasets published from different expeditions (listed on separate rows). For joint expeditions, both contractor codes are listed (e.g. YUZH **/** IOM). ‘Total Cruises’ = total cruises per year, as per available data on the DeepData database. Records from 10 contractors were published on DeepData t the time of the study (July 2021): BGR (Germany), COMRA (China), DORD (Japan), KOREA (Government of the Republic of Korea), GSR (Belgium), IFREMER (France), IOM Interoceanmetal Joint Organisation, OMS (Singapore), UKSRL UK Seabed Resources Limited (United Kingdom), and JSC Yuzhmorgeologiya (YUZH; Russian Federation). There are 16 CCZ-based contractors in total, but 17 contracts (UKSRL holds two separate contracts) and a further two contractors holding licenses outside the CCZ. The six contractors which have active licenses in the CCZ but do not have data published on DeepData at the time of this study include: TOML (Tonga), NORI (Nauru) and MARAWA (Kiribati) (currently all three under an umbrella organisation The Metals Company), CIIC (Cook Islands), CMC (China), and a new contractor, Blue Minerals Jamaica Ltd. It is unclear if the lack of data from these contractors is due entirely to a backlog of data publishing or if not data has been submitted by them.

#### Duplication

We found approximately 6000 duplicate records for the Contractor BGR (Federal Institute for Geosciences and Natural Resources of Germany), and approximately 4000 for UKSRL (UK Seabed Resources Ltd) in the database export. Duplicates were suspected in other Contractor datasets, including KOREA (Government of the Republic of Korea) and IOM (Interoceanmetal Joint Organization), and were confirmed via an OBIS pipeline for identifying duplication in datasets (available in a GitHub notebook, https://iobis.github.io/notebook-duplicates/). We estimate overall duplication is approximately a quarter of the total records assessed (∼10,000 of 40,518). The exact number of duplicates could not be ascertained because of underlying issues with identifiers (detailed in following sections). This duplication appears to have arisen through a combination of issues in versioning of annual contractor data submissions and usage of identifiers. Looking first at versioning, multiple years of the annual data submissions have been published, but in some cases this has resulted in duplicates. ISA are publishing the annual contractor data submissions year by year, from 2015, the year the environmental data template was introduced, and plan to continue until up-to-date (ISA secretariat, pers. comm.). For the yearly data reports, these are either one-off data submissions, e.g. a standalone dataset for a particular cruise that is not then re-submitted the following year, or are iterative data submissions, where records are added to the previous year’s dataset, and any updates added to the existing ones. The latter applies to the UKSRL and BGR annual data submissions for example. However, they have been handled as separate datasets rather than yearly updates, resulting in duplication.

The duplicates are primarily stemming from issues with identifiers. The database export lacks a record identifier (or primary key) and uses the specimen identifier field ‘SampleID’ to reconcile records (Sheldon Carter, pers. comm.). In theory, any records submitted year on year with the same ID should therefore be matched and associated data updated if changed. For the majority of the contractor data submissions, however, unique SampleID values were either not present, or not unique. This applied to all the records for the subset of 2015-2017 data submissions, apart from two contractors (S File 3). Records missing a SampleID value are allocated one during data processing (Sheldon Carter, pers. comm.). Here the possibility arises for duplication. For example, in the BGR data, where no sample IDs were present, records from the 2015 template were allocated a Sample ID when that dataset was uploaded, then the same records allocated a different Sample ID when the 2016 and 2017 data were uploaded, and therefore appear on DeepData output as separate records, producing duplication.

### 4.3. Data fields in DeepData export

Several fields were included in the database output that are not required. For example backend database names were present, ‘AreaKey’; ‘ClusterID’; and ‘BlockID’, and for the latter two, no data entries were present in any case. While the search was for polymetallic nodule data only, the output included fields for vents and sulphide deposits: including ‘HydrothermalActivity’ and ‘HydrothermalVentAge’; and ‘ExtensionPMSSite’. Additional fields were present for taxonomic information, e.g. ‘Subfamily’, the only sub- or super-taxonomic classification field included. Both the reason for its inclusion and the rules around its usage are unclear, as it has been used not for subfamily names, but rather as a field to capture morphospecies, even though there are two separate fields for recording this in the output: ‘Putative.species.name.or.number’, and ‘Morphotype’. Here the former, Putative.species.name.or.number’ has been replaced by ‘Morphotype’ in the 2021 template (S File 4). This may be why both fields were present in the database output, and no entries recorded for ‘Morphotype’.

### 4.4. The ISA Environmental Data Template

The structure of the 2022 environmental data template is split into separate tables by tab, e.g. ‘Point Sample’, ‘Towed Gear Sample’, ‘Chem_Results’, and ‘Biological_Results’. The previous template (2018) was structured with all the tabs (sub-tables) as one wide table. The restructuring into several tables has improved usability, but has also introduced new issues, for example the separation of ‘Point Sample’ and ‘Towed Gear Sample’ tables. The separation of point and trawl data has been made to link biological with resource data in the database as it reflects the underlying structure in the database (ISA secretariat, pers. comm.) but creates an extra processing step that should not be necessary, particularly since sampling information is recorded in a specific field. Also the separation of point and trawl data is not complete, the vast majority of ‘Trawl Line’ data is in fact included in the ‘Point’ data. Examining data fields in the tab ‘Biological_Results’, the 2022 template now includes scientific name; and taxonomic identification qualifier, essential fields for capturing taxonomic identification. These are notable improvements, saving significant processing time. Other key fields were still absent, however, such as a record identifier field that is persistent and unique (equivalent to occurrenceID in DwC; see List of Terms) and as distinct from a specimen identifier, i.e. SampleID (equivalent to catalogNumber in DwC). Another key field missing from the template is an equivalent for the DwC field ‘basisOfRecord’ for designating record type, for example ‘machineObservation’ for an ROV-derived record, or ‘preservedSpecimen’ for a specimen-based one. As in the database output, superfluous fields were present. ‘OrgNum’ for example is a required field (‘TaxaID’ in the previous template) but is an arbitrary number to provide a composite key for ISA data processing. It is therefore a backend column name and as such a redundant field that doesn’t capture any existing data in contractor datasets. It also necessitates an additional processing step by contractors and has the potential to cause confusion. Subfamily is included, but as indicated earlier, this field is not necessary. For the other tabs within the template, superfluous data fields were also present, e.g. ‘Target latitude’/’Target longitude’ in point/towed gear sample tabs.

As a wider observation, some field naming and accompanying definitions are potentially ambiguous. The field ‘MatrixType’ for examples is to capture material or sample type (‘i.e. biological sample, sediment or water unfiltered’), but usage of ‘Matrix’ rather than more intuitive wording such as ‘sample’ or ‘material’ is potentially confusing. A more critical example is ‘SampleID’, which has been interpreted in a variety of ways by contractors. In some datasets, SampleID was used for a batch of samples, equivalent to a deployment or sampling event ID, rather than for an individual specimen record as intended (S File 4 A, B). The current data template includes the field ‘StationID’ for recording station number but this does not account for multiple samplings at a given station, and the template does not include a deployment or sampling event ID to capture this-see Recommendations-identifiers).

Some contractors do not use the SampleID field at all, but rather other fields, such as ‘voucherCode’. Similarly, the field ‘Morphotype’, which is intended to capture morphospecies names could be misinterpreted, as this term usually refers to megafauna identified solely by imagery, as opposed to other types of temporary names such as Molecular Operational Taxonomic Units (MOTUs), which the field is also supposed to capture. The relevant DwC field is ‘taxonConceptID’ which captures all types of open nomenclature or informal species names (11) would be an ideal field name replacement here. As a wider observation, our overall assessment post-testing the new template and examining contractor data submissions (S File 3, 4) is that issues with usage are likely to continue in the new template without a significant re-working including incorporation on rules for filling out required fields.

### 4.5. The OBIS ISA node and DeepData mapping to Darwin Core

The publishing of DeepData records on the OBIS ISA node necessitated a process of mapping contractor data to DwC terms by the ISA data team (see S File 5). The resulting data was later processed by the OBIS secretariat for publication on the OBIS ISA node, documented in a GitHub notebook (https://github.com/iobis/notebook-deepdata). The data processing was done on the datasets mapped to DwC, in JSON format, on the ISA server, not a DwC archive of the DeepData database output itself (S Figure 2). This process of data mapping to DwC by the ISA has resulted in previously missing fields now being incorporated (e.g. scientificName; occurrenceID and basisOfRecord). The DwC terms have been misinterpreted in some places however and mis-mapping of data template fields to DwC was evident. For example, BasisOfRecord, a key DwC term as above for describing the record type has been populated entirely with text entries ‘taxon’. Mapping to DwC terms overall is incomplete, with DwC terms not being utilised where corresponding data are captured in the template, e.g. INSDC accession numbers could be mapped to the term ‘associatedSequences’ in DwC. Some of these fields would be helpful for tracing records and identifying duplication given the lack of adequate record identifiers, e.g. the DwC term ‘datasetName’ would delineate a particular dataset, such as an annual contractor data submission. In some cases, this misinterpretation has produced incorrect taxonomic information. For example, ‘taxonConceptID’, a DwC field recommended by Horton et al., (11) for recording of the open nomenclature name (or taxonomic concept) in DwC terms (11, https://dwc.tdwg.org/terms/#dwc:taxonConceptID), was incorrectly mapped to ‘taxonRemarks’. This has resulted in very low numbers of morphospecies records in the dataset (Figure 2).

**Figure 2.**
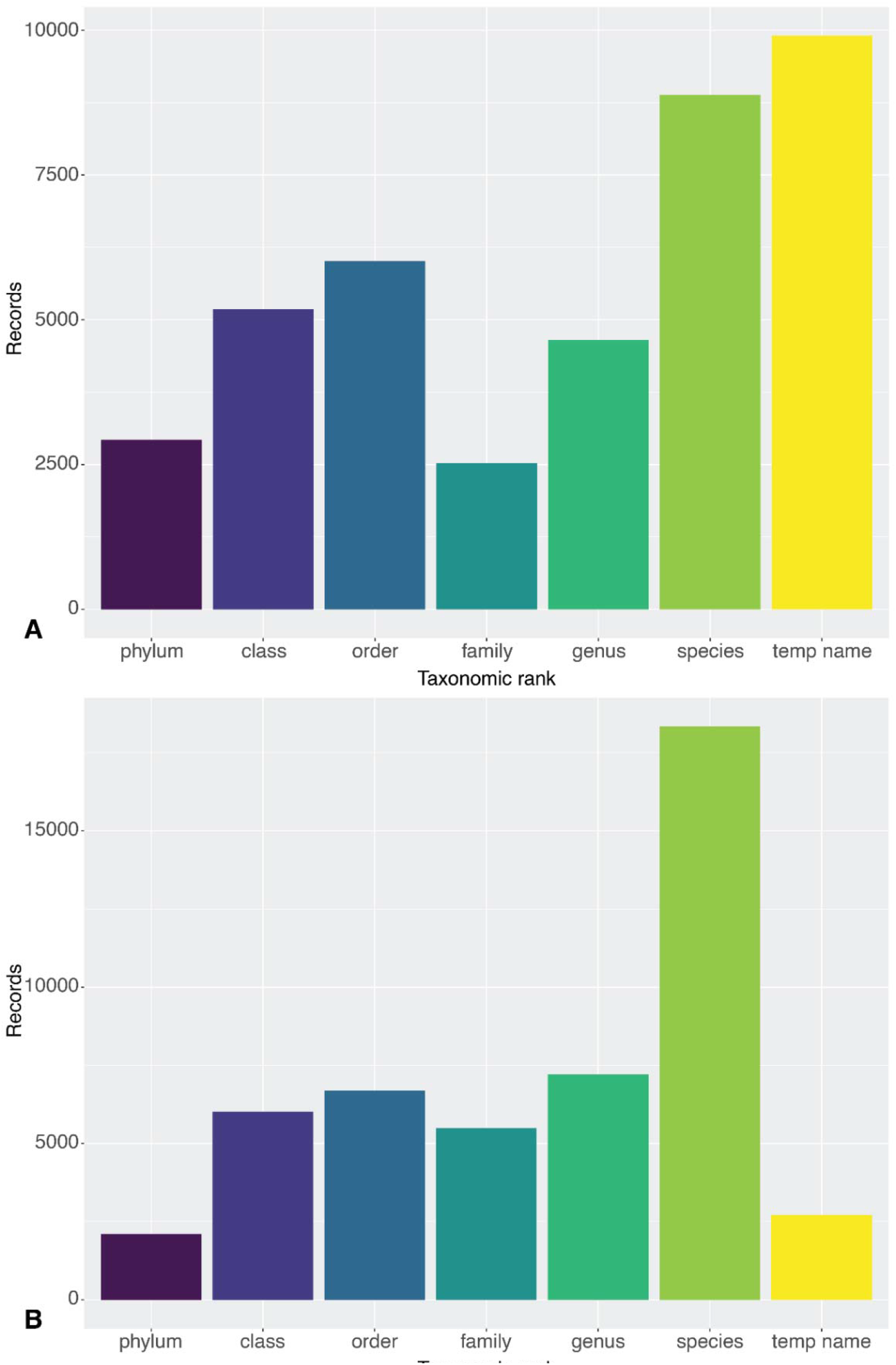
Taxonomic resolution of Clarion-Clipperton Zone DeepData records as published on the 12^th^ of July, 2021, A, from DeepData itself, with 8883 species-level records and almost 9936 temporary name-level records (‘temp name’); and B, via the OBIS ISA node, with very large proportions of records at species level in contrast (18,304) and very few temporary names (2716). Note that temporary names here may include names at levels higher than species-i.e. temporary/informal species names (morphospecies) but also temporary names for higher taxon ranks e.g. undescribed genera and incomplete identifications using open nomenclature.

Additional issues appear to have arisen during data processing for mapping to DwC, also impacting taxonomic information. In the process of mapping to ‘scientificName’ for example, genus names have been duplicated in the resulting scientificName column and the duplicated genus names harvested instead of the species names, resulting in a much lower total number of species names on the OBIS ISA node, 75 compared to the 466 (including pelagic species) from DeepData, as ascertained in a parallel study (Rabone et al., in prep). The duplication of genus name also appears to have resulted in species names being reallocated to other phyla in some cases. For example, some records of the nematode *Capsula galeata* Bussau, 1993 were assigned to the diatom phylum Ocrophyta in the data mapping, presumably because scientificName was designated in DeepData as the genus name only, i.e. ‘*Capsula*’ rather than ‘*Capsula galataea*’; returning *Capsula* J. Brun, 1896 †, an unassigned name in WoRMS. Issues have also arisen in mapping the DwC term for taxonomic rank (‘taxonRank’), where 18,304 records were listed as species, but most were not species level records but at higher taxonomic level, such as genus or family (Figure 2).

Significant issues were also present in treatment of identifiers in the DwC mapping. The DwC term ‘occurrenceID’ is a key, and required field for a persistent, unique record identifier. Here occurrenceID has been generated as a composite key, from combining ‘StationID’/’TrawlID’ and SampleID’. There were duplicates present in this composite key, however. These duplicates were identified by the OBIS secretariat and at the start of the OBIS processing pipeline records were allocated a separate unique identifier. Because of these duplicates in occurrenceID in the DeepData records, a proportion of records cannot be definitively matched between the two databases. Also, the occurrenceID as a non-unique composite key is not present in the DeepData output, only in the JSON files mapped to DwC (and therefore in the OBIS ISA node records), and the composite key would therefore need to be generated with the same formatting to allow any cross-referencing between the records from DeepData or OBIS, i.e. there is not a common record identifier. Even adding the composite key and comparing the records, they do not match of course because the identifier is not unique (and there is different data processing for the DeepData output versus the records on the OBIS ISA node). Overall the number of records for benthic metazoans were different, 40,518 on DeepData and 48,554 in OBIS, which appears to be due in part to slightly more datasets published on OBIS than DeepData at the time of download, but this could not be clearly ascertained because of the underlying identifier issue. In conclusion, standardisation of data to DwC terms to prepare the DeepData records so they can be harvested by OBIS has been a significant step forward, but incorrect data mapping in the process has also compromised data quality.

## 5. Recommendations

The ISA has met a significant challenge to reconcile and publish often variable datasets from contractor annual environmental data submissions. It is a notable achievement that significant biological data holdings (>50,000 records) are now published and available on the database. The 2022 template is also an improvement on the previous version. Through publishing of DeepData records on OBIS, and in the process, mapping data to DwC, some key issues have been addressed and the biological data can now, in part, be classified as FAIR (although reusability is compromised). Despite the issues detailed here, DeepData is a major step forward in developing a centralised repository of biodiversity data in ABNJ, and, given that there has only been four years of development since public release, it is already of great potential value in developing local and regional environmental management plans for this region, and others of our planet that are undergoing rapid industrial exploration.

In a separate study, we have made the first attempt to survey all metazoan biodiversity data from the CCZ using DeepData and published species records (Rabone et al., in prep).

These kinds of regional syntheses would not be possible without the significant efforts from the ISA DeepData team. DeepData provides a crucial source of ‘raw’ occurrence data that are rarely available in publications, even as supplementary files, as revealed in the parallel study. A broader point is that the timing of this work has coincided with a phase of rapid evolution of the database, and that the Secretariat is aware of the limitations discussed here are actively working to address them (ISA Secretariat, pers. comm). There are significant improvements to be made, however, that can address the key data quality issues, with the result of greater utility of the data. It is important to note here that the scope of our study is limited to biological data in the CCZ. Many other data types such as geochemistry data are collected by contractors and held by the database. The FAIRness of these data should also be assessed in depth, especially given these data are only available through DeepData itself, and not also as Darwin Core published on OBIS. Geological data being confidential may be a more complex case, but the potential for greater transparency could be explored as this would have significant scope for improved understanding of ecosystems in the region. Here we provide key recommendations with the aim of improving data quality for both research and environmental policy. These recommendations are also depicted as a potential workflow in Figure 3 and summarised in Table 2.

**Table 2:**
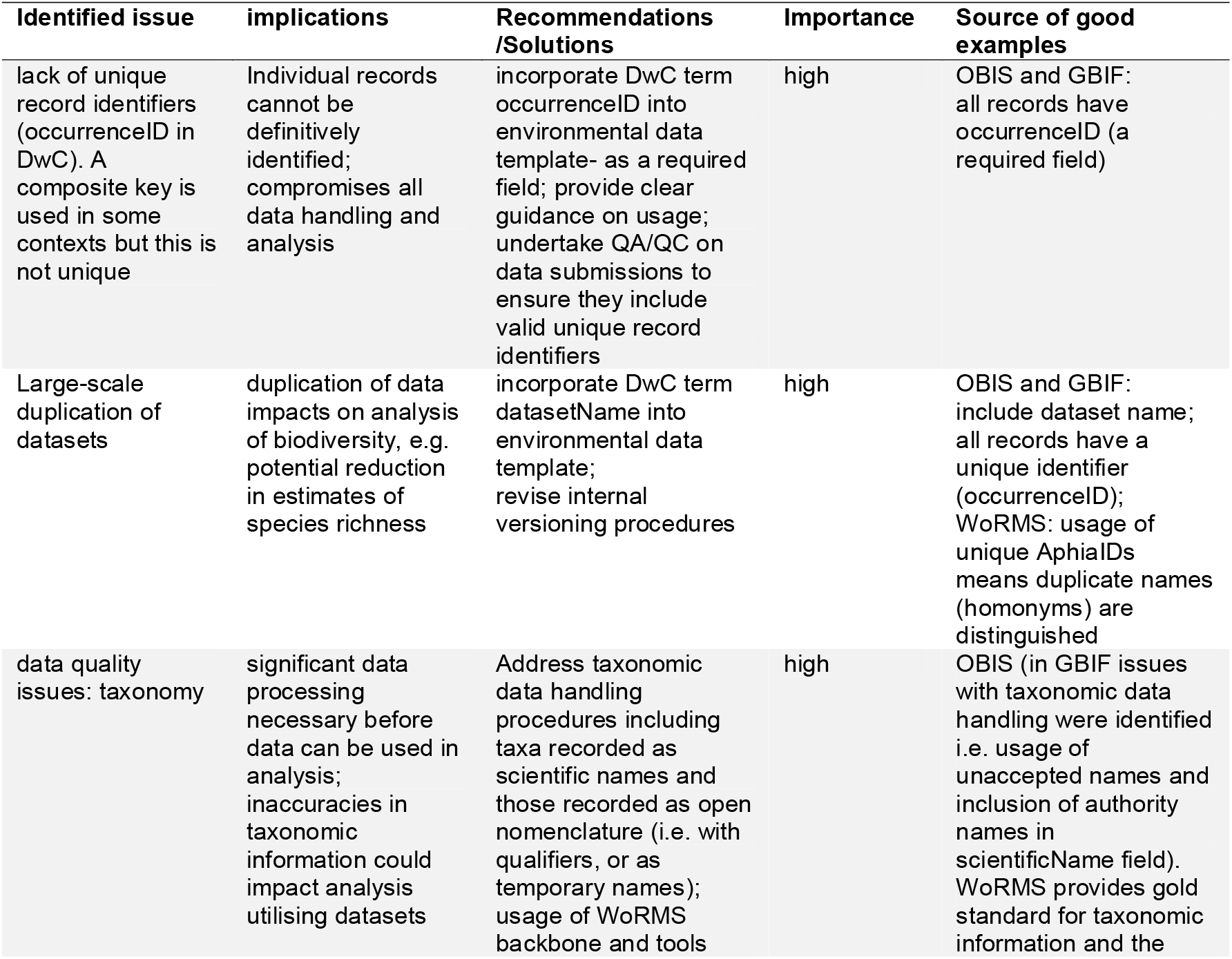
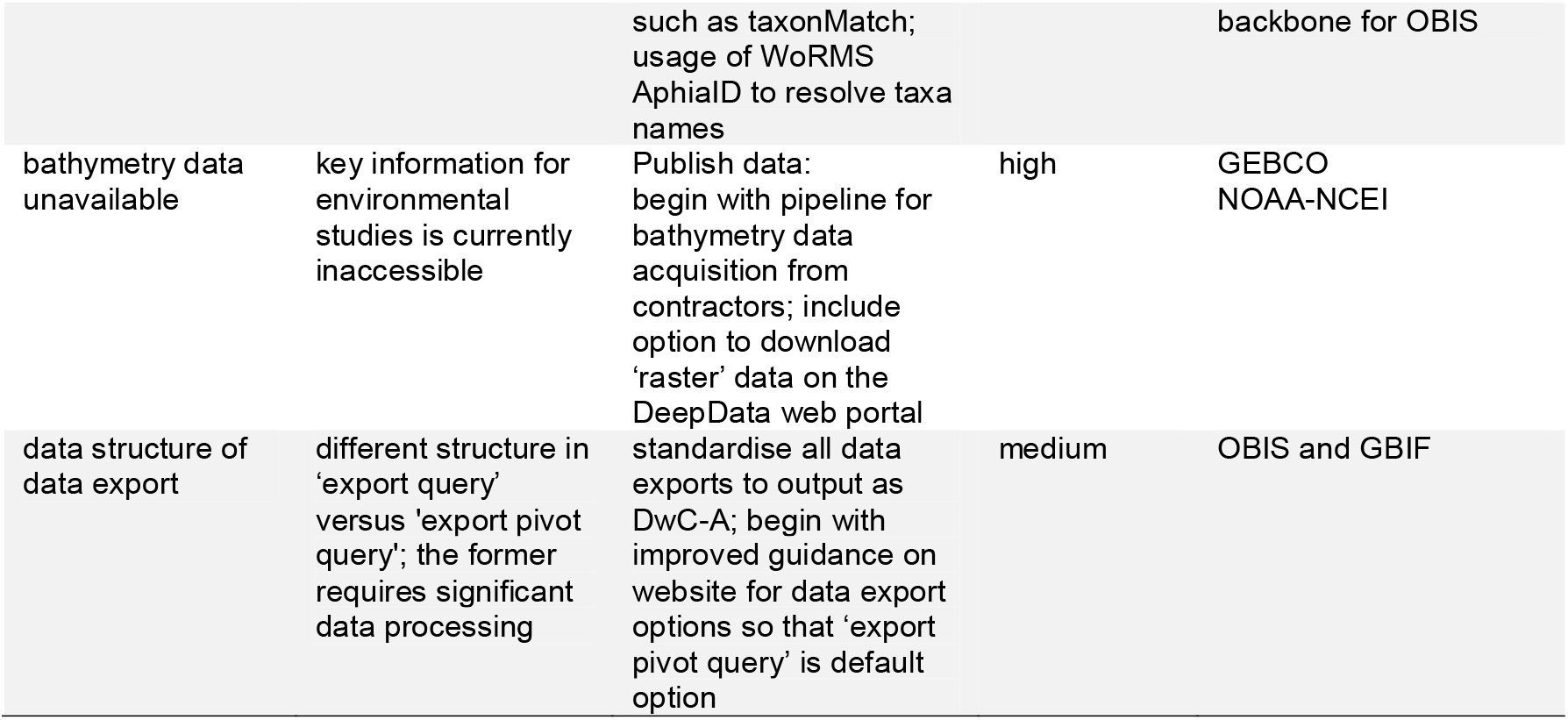
Summary of DeepData assessment, including key current limitations, their implications, and suggested solutions or recommendations to address these, and source of good examples in global databases

**Figure 3.**
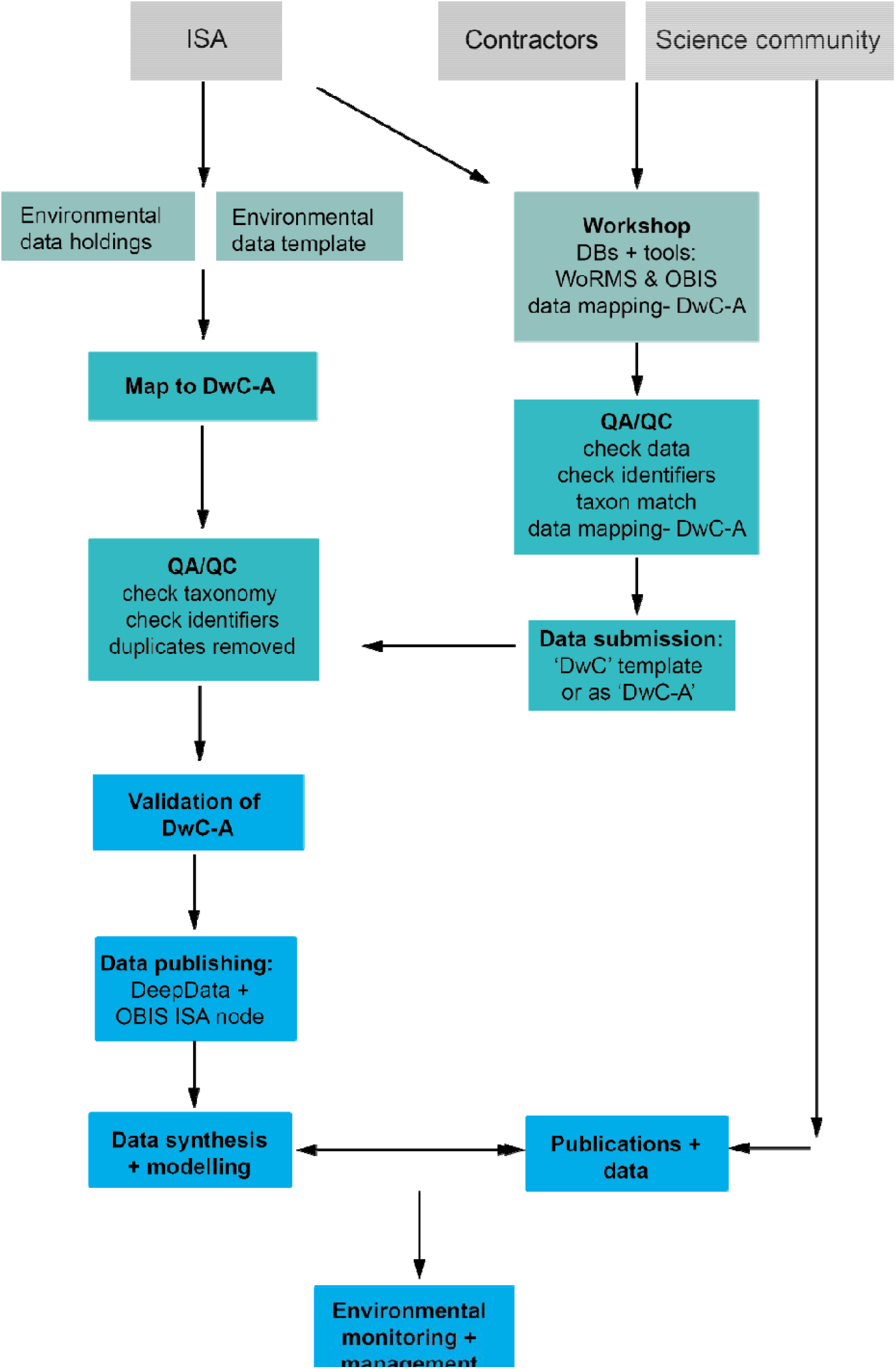
A proposed data management workflow for the ISA. Firstly, the current environmental contractor data submission template is replaced with a DwC compliant version with all fields (column headings) in DwC format, and contractors/data providers can alternatively submit data as a Darwin Core archive file (DwC-A). Existing environmental data holdings are remapped comprehensively to DwC terms as a batch process and undergo QA/QC prior to publication. Concurrently, a public workshop is delivered by the ISA with input from the contractors, the science community and other stakeholders (with full documentation available), covering Darwin Core, databases, in particular WoRMS and OBIS, and tools such as taxon match in WoRMS and tools such as the GBIF Darwin Core validator and assistant. Here contractors undertake QA/QC checks and submit new data (in the new DwC compliant template or as DwC-A). Post QA/QC, dataset are published (as DwC-A) on both DeepData and OBIS on the ISA node. These data subsequently can be utilised for data synthesis and modelling and environmental policy applications

### 5.1. Environmental Data Template column headings replaced with DwC terms, re-mapping of all data to DwC

We make a key recommendation that the ISA update the current environmental contractor data submission template with a DwC compliant version, with all fields (column headings) in DwC format. Darwin Core is a global, community-led, well-established data standard in wide usage and the DwC terms are clearly understood, with a readily available, easy-to-read reference guide (https://dwc.tdwg.org/terms/). To accompany this, we recommend that rules are also incorporated into the template to ensure required fields (e.g. occurrenceID) are populated. Contractors or other stakeholders should also be able to submit data as a DwC archive (DwC-A). The ISA could consider that at a later stage the environmental data template is entirely phased out for a requirement of data submission as DwC-A, i.e. as is the case for OBIS and GBIF. We acknowledge the environmental data template is much broader than the biological data covered here, but data standards including within DwC are available to cover the relevant fields, for example the OBIS-ENV-DATA environmental DwC extension (40). In time, usage of data standards could also be applied to geological data. Full utilisation of the global standard DwC would benefit both the contractors and the ISA data team, as well as other stakeholders and the user community, and would address the key issues we have identified with the database, as outlined here:

- All the fields included in the biological data template can be mapped to DwC terms with less ambiguity and more precision. As a result, data will adhere to a common global data standard, allowing data to meet criteria of being FAIR.
- Essential terms for taxonomic identification that are currently absent from the current database export, e.g. scientificName and identificationQualifier, and other critical required fields such as occurrenceID and basisOfRecord would be included as a matter of course.
- DwC includes the terms ‘verbatimScientificName’ and acceptedScientificName’, therefore the verbatim name as recorded by the contractor, and the accepted name as according to WoRMS identified during data validation (if different) could again be included as a matter of course, which would allow data capture of taxonomic versioning.
- This would significantly reduce the risk of duplication, as the unique identifier occurrenceID is allocated by the contractor, avoiding issues downstream. ISA allocating identifiers as is currently happening is a major breakpoint in the system. Inclusion of the DwC term ‘datasetName’ in the template and proper versioning of annual submissions would further reduce duplication (see the following section 5.2 ‘Data Management Considerations’).
- If data mapping to DwC is done by contractors rather than the ISA, this reduces the possibility for misinterpretation of the data. With adequate training, contractors will be well-equipped to map to DwC terms including those currently misinterpreted by the Secretariat in mapping, e.g. taxonConceptID and basisOfRecord (see following section 5.3 ‘Consultation and training workshops’).
- The data export from DeepData and the OBIS ISA node would be identical. At present, these datasets should contain identical information, but differ owing to the different data processing steps, and more critically, because a unique record identifier is absent from the DeepData export, the records cannot be definitively matched. One the DwC mapping is revised, datasets could be republished on both databases as matching record sets (see section 5.2)
- The DeepData output as a DwC would be FAIR and analysis ready. The current data export from the database requires significant general data processing and even interpretation, for example cleaning of taxonomic information. Similarly significant processing was also required for data downloaded via the OBIS ISA node because of mis-mapping to DwC terms. With correct implementation and interpretation of DwC terms according to established guidelines, in combination with adjustments to ISA workflows as detailed in the following section, the output from both DeepData and the OBIS ISA node would be ready for analysis.
- The database export could be downloaded as a DwC archive (DwC-A; or as a csv file with an xml metadata file). This would standardise the database output structure and allow for proper metadata recording. Currently there are two options for export of biological data: ‘export query’ or ‘export pivot query’. These two options have a different structure (the former requires restructure prior to analysis, as described in results). Full utilisation of DwC would allow for interoperability of the data as the data export could be provided as DwC-A (as currently done for OBIS and GBIF)..
- Making the template fully Darwin Core compliant would allow the ISA to implement the data processing steps currently done by the OBIS secretariat. It could also facilitate the potential automation of the whole submission process and initial QA/QC steps at a later phase of the database

### 5.2. Data Management Considerations

#### Darwin Core and usage of identifiers

We also recommend some key adjustments to data management in the following section to complement the process of address the issues identified and facilitate republishing of these data. Firstly, fully utilising DwC would also necessitate a revision in usage of identifiers (Figure 4). Having datasets with valid unique identifiers is essential and would greatly reduce or even remove duplication. Currently there is no requirement for a record identifier (occurrenceID in DwC) or one present in DeepData (persistent/unique or otherwise) and it is crucial to address this. In DeepData, the specimen identifier SampleID is used as the record identifier (including within a composite key to generate a unique identifier for DwC mapping for harvesting of data by OBIS). This is problematic for several reasons. First, this identifier is often missing from contractor data submissions, or is not unique. Second, given that not all environmental data submissions will be individual specimen-level records, it is not appropriate to utilise it as a proxy ‘universal’ record identifier. Third, good data practice requires that any digital record should have its own unique identifier as a matter of course, as this is crucial to any data handling. In fact, occurrenceID is the sole required field in a DwC data submission to OBIS or GBIF. It should be a unique and meet criteria of persistence, resolvability, discoverability and authority, for example a globally unique identifier or GUID (25, 41, https://dwc.tdwg.org/terms/#dwc:occurrenceID). For examples of usage in the CCZ, see Wiklund et al. (42). ISA allocating identifiers is a key fragility in the system-allocation of unique IDs by contractors would avoid many problems and also mean that the Secretariat would not have to generate a composite key.

**Figure 4.**
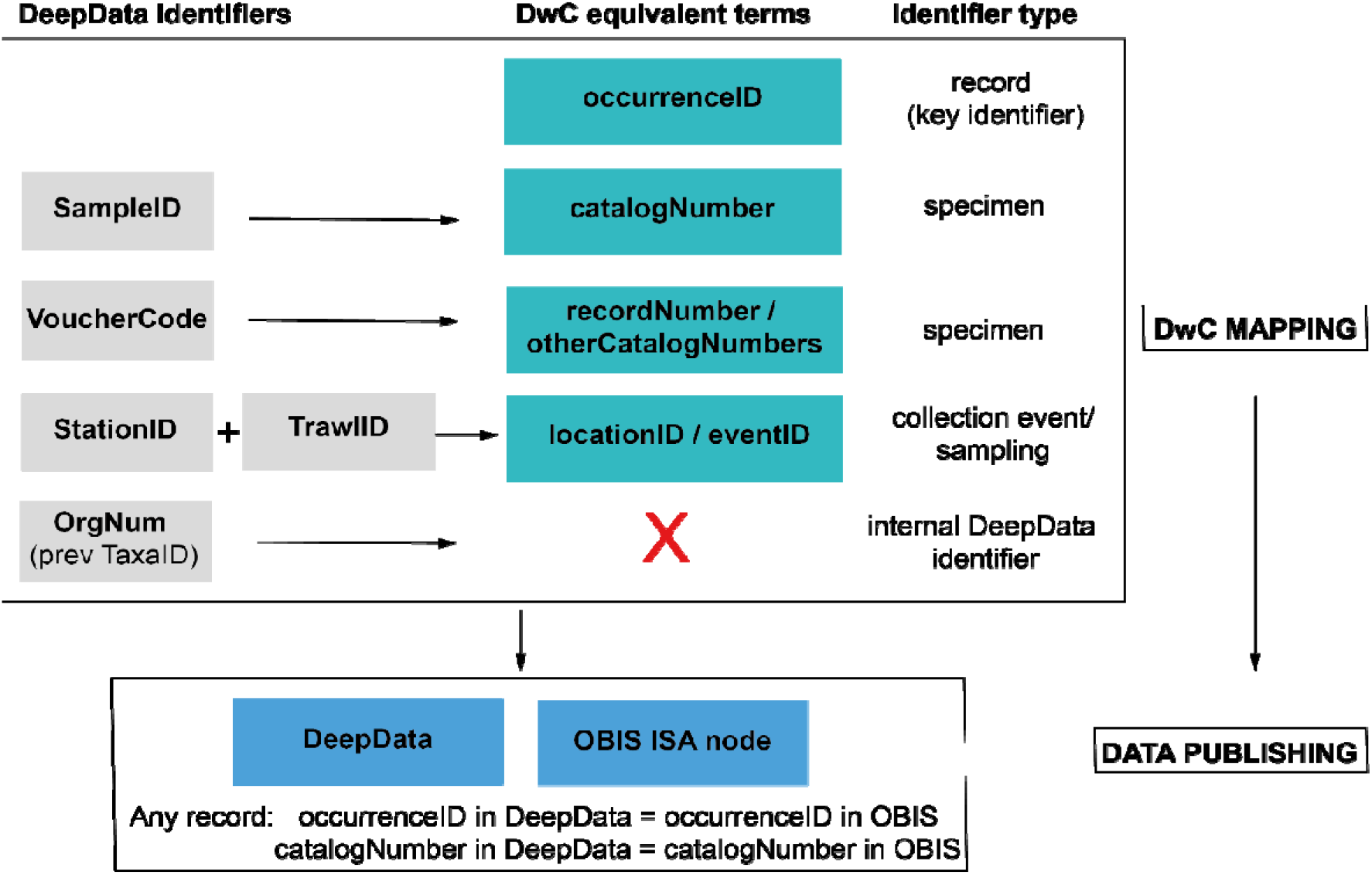
Identifier fields in DeepData, and recommended revision of usage and mapping to equivalent DwC terms. Currently there is no unique record identifier (occurrenceID) in DeepData, or a requirement to include one in the current environmental data template. This key identifier is needed for the data template, database export, and within the database itself. SampleID is currently the key identifier in DeepData and used as a proxy record identifier (although it is neither unique or persistent) and currently mapped to occurrenceID (as in the ISA DwC guidance), but catalogNumber is in fact the equivalent DwC term. VoucherCode instead is currently mapped to catalogNumber, but would be correctly mapped to recordNumber (or otherCatalogNumber). Many other DeepData fields not shown could be better mapped to Darwin Core terms with more precision, for example ‘Morphotype’ replaced with ‘taxonConceptID’. See Recommendations section, ‘Data Management Considerations’ paragraph for details.

Separate specimen identifiers (catalogNumber) are not required in OBIS or GBIF but can support traceability of physical specimens within an institute. The same identifier (i.e. occurrenceID) may be used by some institutes as catalogNumber. However, many collection institutes have a different code (sometimes human readable) and these are used as an internal institutional identifier including on physical specimen labels (Rabone et al., in prep; 43). In the ISA DwC mapping guidance, SampleID has been mapped to occurrenceID, and VoucherCode has been mapped to catalogNumber (S File 5A, B) but this is a misinterpretation of the terms, rather SampleID maps to catalogNumber, and VoucherCode could be mapped to either DwC term otherCatalogNumber or recordNumber (Figure 4).

Usage of sampling event and location identifiers could also be revised. The DeepData sampling event identifiers StationID and TrawlID are included in the template, but these do not allow for recording of different deployments/samplings within a station for example. This is a non-trivial issue as accurate delineation of samples is key in biodiversity analyses. Here the DwC terms locationID and eventID could be utilised (Figure 4). An additional identifier that could be included in the database export is the DwC term associatedSequences to capture INSDC accession numbers, unique identifiers within the INSDC system. For data publishing, revising existing usage of identifiers in DeepData, in particular incorporating occurrenceID would allow records in DeepData and OBIS to be reconciled: any given record in DeepData would have a persistent record identifier-occurrenceID and the corresponding record in OBIS would have the same occurrenceID (Figure 4). Similarly, catalogNumber for the same record in DeepData if present would match the corresponding OBIS record (as would all data fields). Given the centrality of identifiers in data handling, datasets missing unique record identifiers, and specimen identifiers where applicable (i.e. occurrenceID and catalogNumber) would be sent back to the data provider/contractor for revision. Guidelines on best practice in usage of identifiers (23, 24) could be provided by the Secretariat and included in the workshop (section 5.2).

#### Revision of data mapping to Darwin Core

To accompany this process we recommend comprehensive field (re)mapping to DwC for the template and existing data holdings, both data submissions in template-form, and legacy – or pre-template data. The existing DwC data mapping is incomplete and incorrect in some cases (e.g. morphospecies names mapped to taxonRemarks rather than taxonConceptID). It is important to note that because of the current mis-handling of taxonomic data, unsupported scientific conclusions could be drawn without full cleaning and interrogation of the data. More comprehensive mapping will also result in better data capture. For example, some contractors have included non-specimen records, such as image only records in the datasets, which could be described using the basisOfRecord field. While a key to mapping template column headings to DwC is provided, this is somewhat buried in the guidance. This documentation could be revised once the mapping is revised. Data mapping to DwC would also allow for publishing of legacy datasets. This is particularly important given the lack of legacy data available, with very few published works available prior to 2000, as ascertained in the parallel study (Rabone et al., in prep). Although data quality can be highly variable in legacy data, here DeepData could draw on lessons from natural history collections, publishing data with data quality/data completeness flags as done in GBIF for example. The remapping could be done as a batch process with reference to DwC guidance and in consultation with OBIS so that datasets are treated consistently. Adjustments to the data processing pipeline may also be required to avoid taxonomic mismatches such as in the *Monticellina* example detailed in results. This could be achieved by additional scripting in the case of ambiguous taxonomic matches, e.g. where a name matches more than one in the WoRMS database, the higher taxonomy levels are interrogated and cross-referenced.

##### Address duplication in records in DeepData

It is important for the secretariat to prioritise removal of duplicate records in the database as this can impact diversity estimates in any usage of the datasets. Analysis from a parallel study has shown that the duplicates result in reduced diversity estimates (Rabone et al., in prep). As above, there is the possibility of erroneous scientific conclusions if the datasets are used in secondary analysis in their current state. OBIS has provided a pipeline for identifying duplicates, which is a useful tool, but it was not comprehensive in its assessment. Therefore, it is important to make changes to data management, both in usage of identifiers as above, but also at the dataset level. As above, the DwC term datasetName would ideally be included in the template as a required field. Improved versioning and documentation of datasets will assist in both preventing and identifying duplication. Communication and involvement of the contractors will also facilitate this process. Contractors could also be required to do iterative data reporting rather than one-off submissions where applicable, i.e. every year the entire dataset, along with any additional new records are submitted, and no ‘one-off’ data submissions are made. This would ensure that year on year changes to records are captured e.g., updates to taxonomic identifications, and potential for harvesting of duplicate datasets is minimised. We recommend that changes are also made to the ISA data publishing strategy, so that rather than publishing contractor data received from 2015 up to the present, the reverse is applied, i.e., the latest data submissions-post QA/QC are published. Any additional data that are identified from previous years submissions not included in the current submissions, e.g. contracts that are no longer active, are then added. This will again reduce potential duplication. Further, once record identifiers are incorporated into the template itself, i.e. occurrenceID (Figure 4), any duplicates at the record level could be automatically flagged for example through cross-referencing of these identifiers during the submission process.

### 5.3. Consultation and training workshops with contractors and the scientific community

To support the DwC submission process, training and workshops for contractors, also involving the scientific community and other stakeholders could be considered by the ISA. Wider involvement of the scientific community is important, both for user feedback on the database, and to broaden the data-provider base and encourage publication of non-contractor data on DeepData. The workshops could focus on the relevant databases, tools and data standards: in particular, OBIS, WoRMS and DwC. There are also online tools available which could be utilised in the workshop. These include the WoRMS taxon match tool to help with taxonomic data validation, the GBIF Darwin Core assistant and validator, and the Integrated Publishing Toolkit (IPT, 44) to support mapping datasets to DwC.

As missing information in the database is often a result of incomplete contractor data submissions, this could be addressed in a combination of training, consultation with the contractors, documentation and by incorporating rules in the template so that mandatory fields (e.g. occurrenceID) have to be (correctly) populated to submit the data. Key information for biological/ecological studies was often absent from datasets, for example relative density and abundance data; depth, sampling method, taxa identification method, habitat (e.g. nodule/sediment/water column) and broader habitat classification (e.g. ‘seamount’/’abyssal plain’/’rocky outcrop’). These are important data both for deep-sea research and for developing environmental policy both for the region and at broader spatial scales. Establishing a line of communication with the contractors could help address some of these data gaps and wider data quality issues. Together with the DwC submission process and additional QA/QC, this could result in greater quality of submitted data to be ingested into the database, with fewer processing steps required, to the benefit of all stakeholders. A general emphasis should be on quality rather than quantity of the data. While issues remain outstanding, the ISA could consider documenting database limitations clearly on their website to inform end users (including policymakers) before they conduct any analyses.

### 5.4. Potential future developments of DeepData

As DeepData reaches a more mature state, further developments of DeepData would be worthwhile. Our review has focussed in main part on data quality of the biological database output, here we turn to web functionality. It should be noted, however, that as web functionality is inherent to general usability and user experience, it is a key element of general database functionality. Also some of the recommendations listed below, in particular provision of bathymetric data will be critical to characterising deep-sea environments, and therefore should not necessarily be regarded as ‘optional extras’ but rather as core development. Extensive testing of the web interface is recommended. With data systems, usability and user testing is more critical than theories as to how the systems may work. The ISA here could draw on the model of ‘agile’ software development with extensive user testing and response to user feedback. (Rabone & Glover, in review, 45). These developments may also require additional funding. There is an argument to be made for increased resourcing of DeepData given the importance, complexity and scale of the database, and its potential as a decision-making tool for environmental management. This reflects a wider issue in resourcing of biodiversity databases where the fragility of the database funding mechanisms belies their key importance in biodiversity research (20).

- Provision of an API (Application Programming Interface) to allow the database to be directly interrogated. This will be a most useful tool for utilising the database.
- Provision of a DOI (Digital Object Identifier) from DeepData to allow citation of datasets, as currently available for OBIS and GBIF. This would also allow for versioning and traceability as well as data citation.
- Move to web-based data submission platform, where the DwC archive is submitted via the website, and automated QA/QC checks are initiated, e.g. files submitted without valid identifiers could generate an error code as is currently done on web forms.
- Provide information on database and data updates, e.g. when the database has been updated and a list of datasets published. This will support FAIRness of data and general transparency (46, 47, 34). This is currently listed on the website as an
- upcoming feature^5^ (i.e. publication of a file catalogue) and should be straightforward to implement. It could also include a list of submitted datasets that are yet to be published, therefore clarifying which contractors are actively collecting data (Table 1).
- Provide a dynamically updated cruise inventory on the database for all cruises that have taken place up to current cruises and potentially those in planning. This could be very simple with research vessel and contractor name/s, with cruise dates (e.g. Table 1), but would be very helpful information for all stakeholders. This could even provide a model for the cruise notification system proposed in the BBNJ draft treaty text (20).
- The functionality to interrogate data by APEI layer – currently any data outside a contract or reserved area is labelled as ‘OA’ (outside area) rather than with the APEI in question. This requires geographic mapping of records (e.g. in R or QGIS) to ascertain the actual record location. The usability of the web interface could be developed further, for example the ability to click on a section of the map, such as a given contract area or APEI, and a summary of available data for that given region is made visible in a side-bar. Such functionality is not duplicating what is present on the OBIS ISA node and is aligned with the GIS-based focus of DeepData.
- Web functionality whereby taxonomic experts can flag erroneous identifications in records on the web portal, as ‘community curation’. This is possible in both WoRMS and GBIF (via different mechanisms, e.g. in WoRMS, taxonomic editors can add or edit records). As similar functionality is planned in OBIS, when this feature is live in OBIS, potentially the ISA data team could be alerted to any tagged records via the OBIS ISA node. This could allow for simple errors, such as pelagic species recorded as benthic, to be identified. As a wider point, a pipeline to identify pelagic taxa recorded as collected from benthic samples, found to be extensive in DeepData (Rabone & Glover, in review, Rabone et al., in prep) would be of great benefit and could be considered as an additional taxonomy QA/QC step, for example the cleaned taxa names compared to attribute data in WoRMS and pelagic species named could be tagged.
- Development of a data dashboard on DeepData for interrogating, summarising and visualising the data. The emphasis should be on making the dashboard as simple as possible. Now that data is available on OBIS on the ISA node, where there is significant functionality for summarising and visualising data, there may not be the same imperative to develop the web interface. However, different databases have different user communities; and some stakeholders are likely only to use DeepData and not the OBIS ISA node. This dashboard may be particularly helpful for policymakers, who may be less likely to download and analyse the database holdings. It will also support FAIRness of the data and general transparency.

An additional improvement for DeepData could include the storage of relevant literature, straightforward given existing functionality to store documentation (‘Docs’ tab on the website; S Figure 1). Our parallel study shows some key data gaps in DeepData where information is available in the literature (Rabone et al., in prep). This would also align with the ISA mandate to facilitate and support marine scientific research in the Area. Similarly. DeepData could include storage and handling of image data, for example megafauna specimen imagery or in-situ seabed images. Again with the ‘Photo’/’Video Gallery’ tab in DeepData, the functionality to store and publish imagery is already in place. There is a precedent here also with the CCFZ image atlas for in situ imagery, “Atlas of Abyssal Megafauna Morphotypes of the Clipperton-Clarion Fracture Zone” co-administered by the ISA, which was in wide usage by researchers (e.g. 48). However, image data is computationally expensive in terms of required storage, and the technicalities of handling more complex data types. DeepData could potentially partner with platforms such as Bio-Image Indexing and Graphical Labelling Environment; BIIGLE (49), to provide images with metadata to develop image libraries. The mechanics of how this partnership could work in practice may need some thought as image annotation platforms like BIIGLE do not tend to specialise in storing imagery-but there is a clear need for such functionality. Databases of imagery with quality metadata could support machine learning identification efforts, as currently done with iNaturalist, a global citizen science application for recording species observations (https://www.inaturalist.org/).

This could be extended to acoustic images e.g. multibeam imagery for bathymetry. Bathymetric data is listed on the database but not available other than a small amount of bathymetry metadata for a sole contractor. Given the categorisation of data into ‘point’ and ‘line’ on the database, a category for ‘raster’ or similar should be added to allow for bathymetry data which is typically in this format. Bathymetry are the first datasets collected in deep-sea surveys, and essential to ecological studies. For DeepData to fulfil criteria of FAIR, it is important for these data to be made available, here the ISA should make the provision of bathymetry from all offshore campaigns a requirement, and to develop a pipeline for publication of these data. Here DeepData could also work directly with the Multibeam Bathymetry Database supported by the National Oceanic and Atmospheric Administration (NOAA^6^, https://doi.org/doi:10.7289/V56T0JNC), or the GEBCO-Nippon Foundation SeaBed 2030 project (50); where, as for WoRMS and OBIS, existing partnerships are in place.

## 6. Concluding Remarks

While our study is focussed on the CCZ and the ISA database, it illustrates some of the challenges – and opportunities for biodiversity databases, in particular to improve their utility for both research and environmental policy. The DeepData collaborations with OBIS and WoRMS and data mapping to Darwin Core heralds a welcome new phase and rapid evolution for the database and ISA data management practices. DeepData is now integrating with global databases, and global common data standards, allowing for data exchange and integration, for data to be FAIR. However, notable, non-trivial issues with data quality remain, particularly regarding identifiers, duplication, and treatment of taxonomic information. Our review of the database has illustrated the integral importance of global community-led data standards and persistent identifiers for biodiversity data. While the challenges of DeepData reflect those in the wider biodiversity data ecosystem, given the direct connection of the database with the regulator, and its potential to be directly utilised in development of environmental policy, it is even more urgent that these issues are addressed. It would be of great value to be able to directly interrogate the database for species distribution or diversity for example, or on a regional scale, DeepData could ultimately become critical in helping to develop the REMP for the CCZ and other seabed regions managed by the ISA. There is the potential for DeepData to provide an invaluable resource both for research and environmental management. The database is at a nascent phase of its development, here engagement and involvement of the science community, policymakers and contractors to further the development of DeepData is obviously critical. While feedback from user communities of databases via feature requests or bug tracking for example is common practice, more formal and comprehensive assessments of databases like the current study are rare, and we hope in the process to have provided the ISA with useful and implementable recommendations. There is a collective responsibility amongst all stakeholders to support open data efforts such as DeepData and community data curation. However, the ISA is well placed to lead and coordinate activities and encourage efforts in best practice and eventually may even provide an exemplar for high quality deep-sea biological datasets. Such information could be utilised for biodiversity assessments and observing programmes, including contributions to indicators and variables such as EOVs and EBVs (3, 4). These could be applied at regional scales with DeepData contributing information to the proposed Deep Ocean Observing Strategy DOOS demonstration project for the CCZ (51); and even at global scales, across ocean basins. In time, DeepData may be viewed not through a CCZ or even an ISA lens but rather through a global one and as part of the global biodiversity data landscape. An ultimate focus on the importance of biodiversity data to support conservation efforts is key. Partnerships with big international science programs including the UN Decade of Ocean Science, DOOS, major genomic data projects like Earth Biogenomes (52), and the GEBCO Seabed 2030 mapping programme (among others) will be crucial, as will integration into the wider policy landscape, i.e. the UN Biodiversity Beyond National Jurisdictions (BBNJ) treaty process.

## 7. Data Availability Statement

All datasets and code are available as supplementary files. This paper is part of a larger study: Rabone, M., Glover, A.G., 2022, A review and synthesis of CCZ benthic metazoan biodiversity data from the ISA DeepData database, the literature and other published sources. Report prepared for The Pew Charitable Trusts. Report number NHM SON20001/PewFR

## 9. Acknowledgements

Funding for this project was received from The Pew Charitable Trusts to the Natural History Museum. This work involved the first formal collaboration of The Pew Charitable Trusts, the Natural History Museum, and the International Seabed Authority. We are very grateful to Andrew Friedman, Chris Pickens and Peter Edwards of The Pew Charitable Trusts for their support and assistance throughout the project. We would also like to thank Luciana Genio, Sheldon Carter, Tamique Lewis and Ansel Cadien, of the ISA Secretariat for their cooperation and assistance; Pieter Provoost and Ward Appeltans, OBIS Secretariat for assistance and background information on the databases; Hannah Lily (formerly of The Pew Charitable Trusts) and Diva Amon for preliminary discussion on the project, Ani Chakraborty of Pew for project advice and Harry Rousham and Robyn Fryer of NHM for administrative support, and the Nautilus biodiversity data working group where a Miro board inspired elements of Figure 1. MER and AGG are currently funded by the UK Department for Environment, Food and Rural Affairs (DEFRA) Global Centre on Biodiversity for Climate GCBC programme.

## Author Contributions

MER with input from AGG conceived the study and designed the methods approach. MER curated data, conducted investigation and analysis, created figures and wrote the original draft. TH, DOB, ESL and AGG edited and reviewed the manuscript drafts. All authors read and approved the final manuscript.

## 10. Figures

**S Figure 1.**
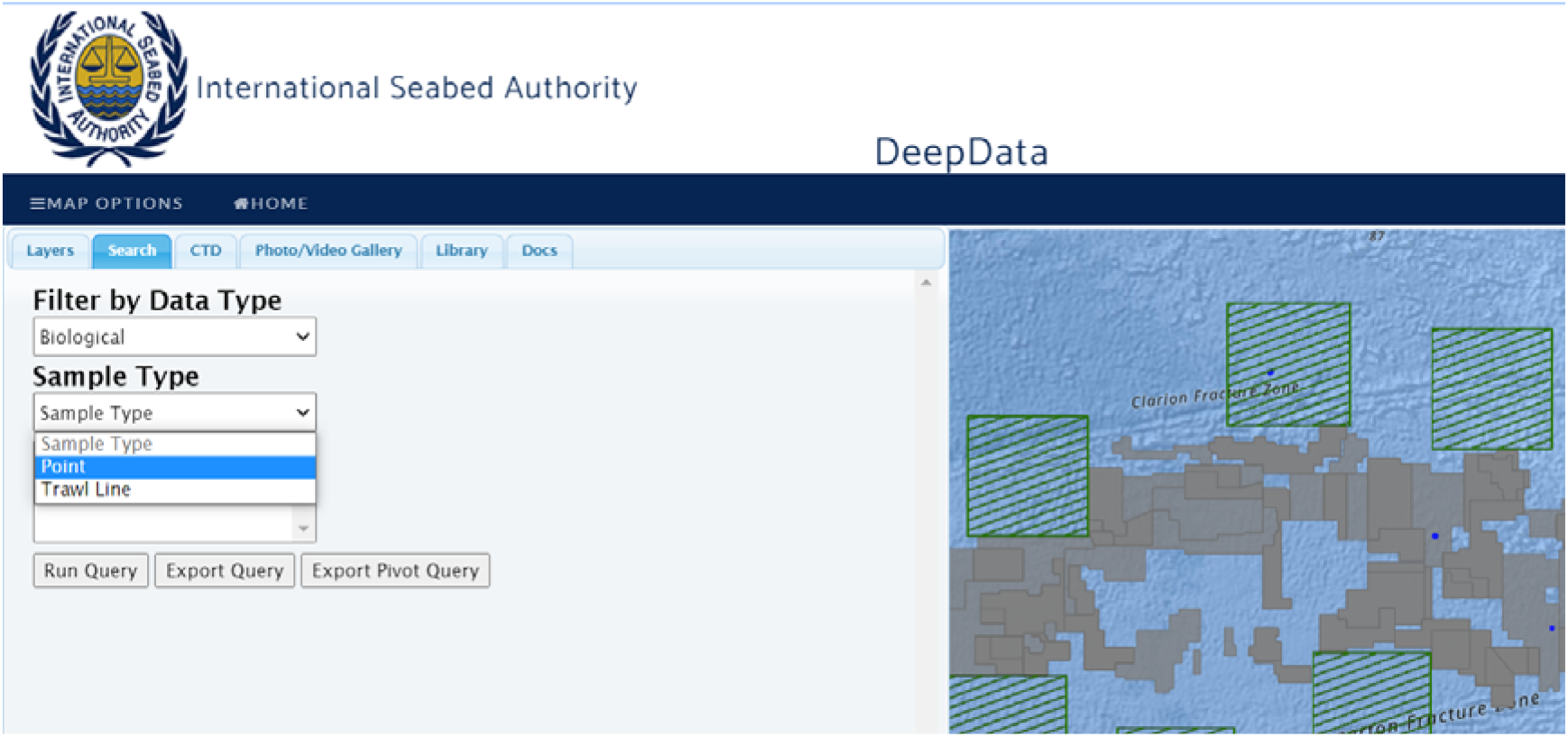
DeepData interface (tab ‘MAP OPTIONS’, adjacent tab ‘MAP’ is a full-page map view) with the main layout of search tabs. View shows the main search tab with data selection menu on the left-hand side and map view on the right (adjacent tab ‘Layers’ with options to pre-select data by contractor and area as described in methods). Shown here, selection for data type ‘Biological’ (the option being ‘Environmental Chemistry), and for sample type ‘Point’ or ‘Trawl Line’ data,

**S Figure 2.**
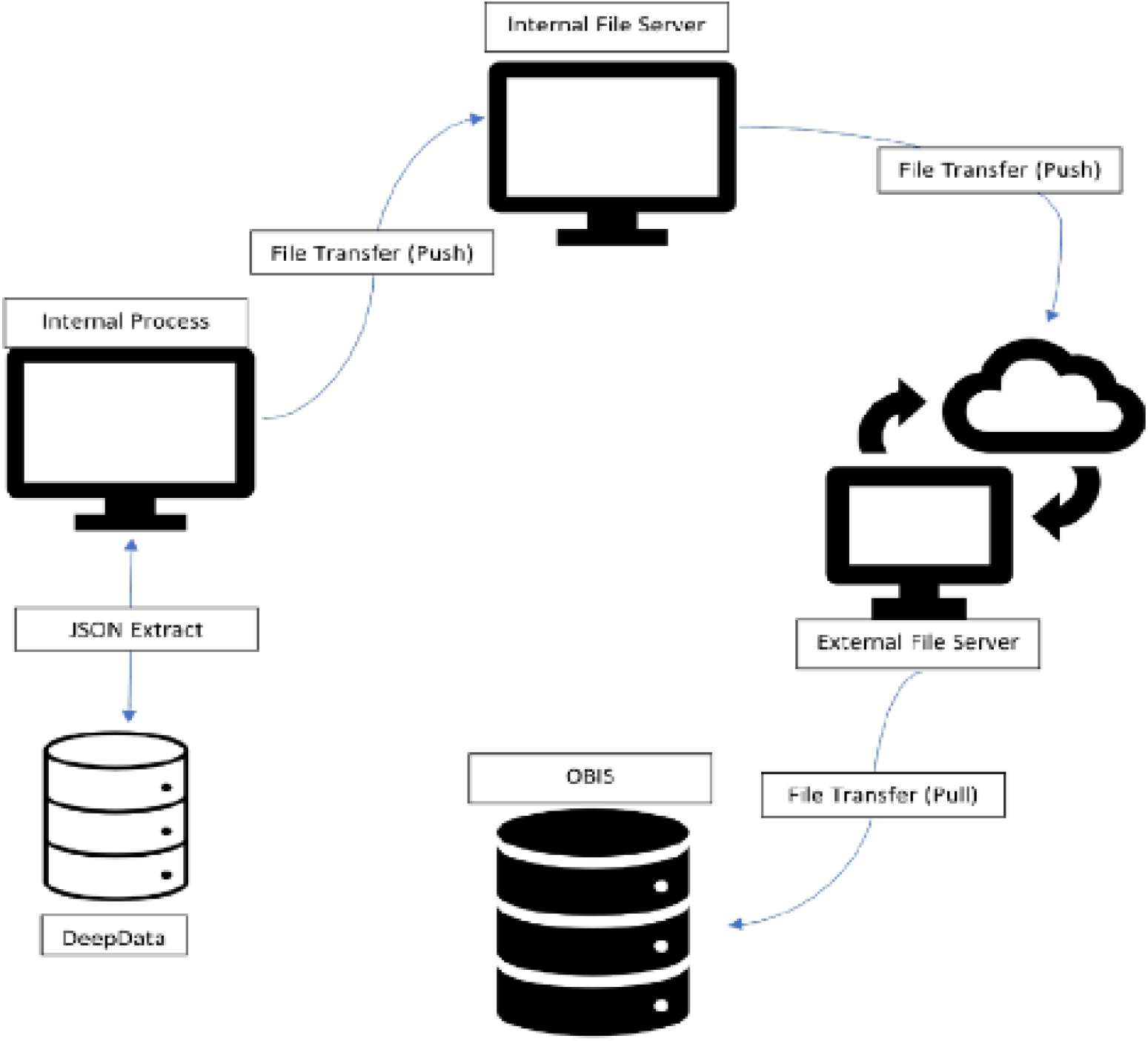
File transfer protocol for datasets from DeepData to OBIS (ISA documentation ‘Deep Data Correlation to DwC in the Context of OBIS. 20th Nov 2020’). Data holdings in DeepData are mapped to Darwin Core, stored in JSON format on an internal server and later harvested by OBIS (from an external server).

## 11. Tables

**S Table 1.** A subset of the DeepData database export file (5 fields of 49), showing how observations are distributed both across rows and columns, or a combination of both wide and long format in the data. Text in bold italic represent distinct observations, text in grey, repeated, redundant data. The ‘Analysis’ field represents column headings (from the data template) and is paired with the adjacent ‘Result’ field (therefore in ‘long’ format, unlike the rest of the data. The data were restructured to entirely wide format- ‘spread’ over separate columns, e.g. for ‘Ecology’ data in the ‘Category’ column, the ‘Nominal size Category’ (in ‘Analysis’) became a separate column (e.g. adjacent to the ‘Result’ column), with the entries as recorded in ‘Result’, i.e. ‘macro’

| ContractorID | Category | TaxaID | Analysis | Result |
| --- | --- | --- | --- | --- |
| BGR1 | Ecology | 612 | Nominal size Category | **macro** |
| BGR1 | Ecology | 612 | Number of individuals | **1** |
| BGR1 | Organism Details | 612 | Sex | **Unidentified** |
| BGR1 | Taxonomist information | 612 | Taxonomist | **Not Reported** |
| BGR1 | Taxonomist information | 612 | Taxonomist E-mail | **Not Reported** |
| BGR1 | Taxonomist information | 612 | Taxonomist Institution | **Not Reported** |
| BGR1 | Taxonomy ID | 612 | Class | **Polychaeta** |
| BGR1 | Taxonomy ID | 612 | Identification Method | **Not Reported** |
| BGR1 | Taxonomy ID | 612 | Kingdom | **Not Reported** |
| BGR1 | Taxonomy ID | 612 | Morphotype | **Not Reported** |
| BGR1 | Taxonomy ID | 612 | Order | **Not Reported** |
| BGR1 | Taxonomy ID | 612 | Phylum | **Annelida** |
| BGR1 | Taxonomy information | 612 | Database Taxa ID | **Not Reported** |
| BGR1 | Taxonomy information | 612 | Taxonomic Database | **Not Reported** |
| BGR1 | Taxonomy information | 612 | Taxonomic Status | **Uncertain** |
| BGR1 | Ecology | 613 | Nominal size Category | **macro** |
| BGR1 | Ecology | 613 | Number of individuals | **1** |
| BGR1 | Organism Details | 613 | Sex | **Unidentified** |
| BGR1 | Taxonomist information | 613 | Taxonomist | **Not Reported** |
| BGR1 | Taxonomist information | 613 | Taxonomist E-mail | **Not Reported** |
| BGR1 | Taxonomist information | 613 | Taxonomist Institution | **Not Reported** |
| BGR1 | Taxonomy ID | 613 | Class | **Polychaeta** |
| BGR1 | Taxonomy ID | 613 | Identification Method | *Not Reported* |
| BGR1 | Taxonomy ID | 613 | Kingdom | *Not Reported* |
| BGR1 | Taxonomy ID | 613 | Morphotype | *Not Reported* |
| BGR1 | Taxonomy ID | 613 | Order | *Not Reported* |

## 12. Supplementary Data

### Supplementary Data File 1A

All biological data ‘Point’ records published on DeepData on 12th of July, 2021 (raw data download archive file)

SDF1A_DeepData_biology_points_export_2022-07-12.zip

### Supplementary Data File 1B

All biological data ‘line’ records published on DeepData on 12th of July, 2021 (raw data download archive file)

SDF1B_DeepData_biology_lines_export_2022-07-12.zip

### Supplementary Data File 1C

All biological data records published on DeepData on 12th of July, 2021, processed File SDF 1C, “SDF1C_DD_PUBLISHED_main_file_2021-07-12_v10.csv”, metadata file “SDF1C_DD_PUBLISHED_main_file_2021-07-12_v10_meta”

### Supplementary Data File 1D

All benthic metazoan biological data records published on DeepData, final for analysis File SDF 1D, “SDF1D_DD_PUBLISHED_4analysis_ed_2022-05-24.csv”, metadata file “SDF1D_DD_PUBLISHED_4analysis_ed_2022-05-24_meta.csv”,

### Supplementary Data File 2

Biological records from the CCZ region within ISA jurisdiction-contract areas, reserved areas or APEIs, published on the OBIS ISA node, 12th of July, 2021, in DwC format.

File SDF 2, “SDF2_OBIS_DD_4_analysis_2022-04-20.csv”, metadata file “SDF2_OBIS_DD_4_analysis_2022-04-20_meta.csv”

### Supplementary Data File 3

Subset of annual Contractor data submissions, from data templates submitted between 2015-2017

File SDF 3 “SDF7_DD_Contractor_data_raw_files_2021-04-02.csv”

### Supplementary File 4

ISA data reporting template-version 9th of June, 2021. File SF 1 “SF1_Env_Template_20181005.xlsx”.

### Supplementary File 4A

ISA data reporting template guidance-version 1.6, 9th of June, 2021. File SF4A “SF4A_ReportingTemplates_Guidance_v1.6_20210609.pdf”

### Supplementary File 5A

ISA Documentation: “Deep Data Correlation to DwC in the Context of OBIS. 20^th^ Nov 2020” File SF5A “SF5A_Deep Data Darwin Core Mapping 2.pdf”

### Supplementary File 5B

ISA Data file: DeepData mapping to Darwin Core File SF5B “SF5B_DeepDataDarwinCoreDump.xlsx”

### Supplementary File 6

R script for data collection, processing and analysis. FILE SF6 “DeepData_review_data_processing_script.R”

## 13. List of Terms and Abbreviations

(terms hyperlinked and in bold-DwC terms)

ABNJ: Areas Beyond National Jurisdiction
APEI: Area of Particular Environmental Interest regions designated by the ISA as potential regional conservation zones
API: Application Programming Interface
AUV: Automated Underwater Vehicle
BasisOfRecord: DwC term to describe record type (‘the specific nature of the data record’) e.g. preservedSpecimen
CCZ: Clarion Clipperton Zone, also known as the Clarion Clipperton Fracture Zone (CCFZ)
Checklist: Inventory of species/taxa names, often organised by taxonomic group or region **<u>catalogNumber</u>** DwC term for specimen identifier (An identifier (preferably unique) for the record within the data set or collection’)
Contractors: holders of mineral exploration contracts
DwC: Darwin Core, a global data standard administered by TDWG (Biodiversity Information Standards, formerly the Taxonomic Databases Working Group).
DOI: Digital Object Identifier
EIA: Environmental Impact Assessment
FAIR: Findable, Accessible, Interoperable and Reusable
GBIF: Global Biodiversity Information Facility
GIS: Geographic Information Systems
GUID: Globally Unique Identifier identification qualifier taxonomic identification qualifiers, such as aff. cf. sp. nov. (**<u>identificationQualifier</u>** in DwC terminology)
INSDC: International Nucleotide Sequence Database Collaboration
ISA: International Seabed Authority
JSON: JavaScript Object Notation
LTC: Legal and Technical Commission
MOTUs: Molecular Operational Taxonomic Units; a type if informal or temporary name (open nomenclature)
Morphospecies: These are informal working species names used prior to formal description, also known as morphotypes, OTUs, working species, or temporary names.
NCBI: National Center for Biotechnology Information (administrates the GenBank database)
OBIS: Ocean Biodiversity Information System **<u>occurrenceID</u>** DwC term for record identifier (‘An identifier for the Occurrence (as opposed to a particular digital record of the occurrence). In the absence of a persistent global unique identifier, construct one from a combination of identifiers in the record that will most closely make the occurrenceID globally unique’)
Occurrence data: distributional records of species/taxa
Open nomenclature: system of signs to describe uncertainty around identifications, or designate informal/temporary taxa names prior to formal description (e.g. morphospecies)
REMP: Regional Environmental Management Plan
ROV: Remotely Operated Vehicle
Scientific name: The designation or identification of an organism (**<u>scientificName</u>** in DwC terminology)
taxonConceptID: DwC term for open nomenclature-temporary/informal names (‘n identifier for the taxonomic concept to which the record refers -not for the nomenclatural details of a taxon’)
TDWG: Biodiversity Information Standards (formerly Taxonomic Databases Working Group)
QA/QC: quality assurance/quality control, here referring to data QA/QC
WoRDSS: World Register of Deep-Sea Species
WoRMS: World Register of Marine Species

## Footnotes

1 https://www.marinespecies.org

2 ISBA/8/C/6; <u>ISBA/5/C/6</u>; <u>ISBA/21/C/16</u>; <u>ISBA/22/LTC/15</u>

3 https://obis.org/node/9d2d95be-32eb-4d81-8911-32cb8bc641c8

4 Here Contractors are listed by mineral type (CRFC, PMN, PMS), with separate entries for the same Contractor holding contracts in different mineral types (https://data.isa.org.jm/isa/map/)

5 For clarity and transparency purposes, the ISA Secretariat will publish a file catalogue on regular basis, listing all publicly available data files contained in DeepData. (https://www.isa.org.jm/deepdata/about#block-seabed-page-title)

6 NOAA National Centers for Environmental Information. 2004: Multibeam Bathymetry Database (MBBDB).

