## Supplementary material for "A review of the International Seabed Authority database DeepData: challenges and opportunities in the UN Ocean Decade": SF4A_ReportingTemplates_Guidance_v1.6_20210609.pdf

Data Management Strategy

### DeepData

### Reporting Template Guidance for the Submission of Digital Data

#### Version 1.6

May 2021

12.2 Details on Identification Methods and Taxonomy Information ... 17
**12.2.1 Morphological ID** ... 17
**12.2.2 Genetic ID** ... 18
**12.2.3 Remote ID** ... 18
12.3 Details on Vouchers ... 19
12.4 Submitting Ancillary Information ... 20
12.5 Provisional Results ... 20
12.6 Ecological and Organism details ... 21
12.7 Chem_Results in the ENV Template ... 21
**METADATA TEMPLATE & DATA SUBMISSION** ... 22
**13. METADATA TEMPLATE** ... 22
13.1 File Naming Convention for Metadata Template ... 22
13.2 Modifying the Template Structure ... 22
13.3 Metadata Template Management ... 22
13.4 Instructions for Populating ... 23
13.5 Photo_Video Theme ... 23
13.6 File Naming Convention for Data Submissions ... 24
13.7 ErrorChecker ... 25
**14. SUBMITTING DATA** ... 25
14.1 Resubmittals ... 25
**15. REFERENCES** ... 26
**Annex A** ... 27
**Annex B** ... 28
**Annex C** ... 29

### Glossary of Terms

**Contractors**
The groups granted a contract by the International Seabed Authority for the exploration of polymetallic nodules, polymetallic sulphides and cobalt-rich ferromanganese crusts.

**CTD**
Pertaining to a system for measuring Conductivity (a measurement of salinity), Temperature and Depth (determined from pressure measurements). Additional parameters, such as pH and dissolved oxygen concentration, can be measured if optional sensors are installed. Often has Niskin bottles attached for taking water samples at specific depths.

**Geodatabase**
A database structure including a spatial component designed to enable mapping.

**Legal and Technical Commission**
The Legal and Technical Commission comprises 22 members elected by the Council of the International Seabed Authority on the basis of their qualifications in the fields of mineral resources, oceanography, protection of the marine environment, or economic or legal matters relating to ocean mining. Members of the Commission are elected for a period of 5 years.

**Metadata**
A set of data that describes and gives information about other data.

**Metadata Template**
Template for recording information associated with all uploaded files.

**Reporting Template**
Templates for submitting structured data.

**Structured data**
Contractor data submittal of required environmental, biological, geological, and chemical data entered in the Reporting Templates.

**Unstructured data**
Contractor data submittal of files and information not accommodated by the Reporting Templates; e.g., photos, videos, maps, calibration or QA/QC data, sidescan/seismic files.

### Reporting Templates – General Instructions

#### 1. Introduction

##### 1.1 Organization of this Guidance Document

This document includes guidance and instructions for populating the Reporting Templates (GEO and ENV) and the Metadata Template. The document has three main parts:

1. **Reporting Templates – General Instructions.** These instructions apply to both the GEO and ENV Reporting Templates (Section 1 – 10).
2. **Reporting Templates – Specific Instructions.** These are specific instructions for entering results into the GEO Reporting Template (Section 11) and the ENV Reporting Template (Section 12)
3. **Metadata Template & Data Submission.** These instructions apply to preparing the Metadata Template and data submission packages to upload to the DeepData website (Section 13 - 14).

##### 1.2 Which Template to Use?

The Reporting Templates are to be used by Contractors and other stakeholders (termed data providers) to submit Structured data (required biological, geological, and environmental chemistry data collected during expeditions) for PMN, CFC, and PMS environments.

Use the GEO_Template for geological results from samples that are considered to represent mineral resources (e.g., geotechnical and minerology properties of PMN, PMS or CFC resources). These results will be flagged as confidential in DeepData (ISA 2015)[1]. See Section 11 for specific instructions on populating the GEO template.

Use the ENV_Template for biological and environmental chemistry results from samples that are not considered to represent mineral resources (e.g., pelagic sediment chemistry, water chemistry, CTD cast sampling). These results will be flagged as publicly available in DeepData (ISA 2015)[1]. See Section 12 for specific instructions on populating the ENV template.

The Reporting Templates can be accessed from the 'Downloads' tab of the DeepData website (https://data.isa.org.jm/isa/map/). A Template Example Package is also available for download and will be an important reference for populating the Templates.

[1] Regulations pertaining to confidentiality of data and information included in ISA (2015):
PMN: ISBA/19/C/17: Part II, Regulation 7; Part VI Regulation 36 and Regulation 37
PMS: ISBA/16/A/12/Rev.1: Part II, Regulation 7; Part VI Regulation 38 and Regulation 39
CFC: ISBA/18/A/11: Part II, Regulation 7; Part VI Regulation 38 and Regulation 39

For questions about how to fill out the Reporting Templates, contact the ISA Data Management Team at:.

#### 1.3 File Name Convention

You may not submit multiple template files with the same file name.

Use the following naming convention for the Reporting Template file:
**ContractID[2] + Contractual Year_+ master file name_ + descriptive details**
For example: BGRPMN12015_Geo_Template_2014rock.xlsm

The file name may not exceed 120 characters.

#### 1.4 Modifying the Template Structure

The Template structure is designed to be compatible with ETL procedures to append results into the DeepData database. As such, the structure is not flexible:

- All fields in these worksheets are text and should remain as text.
- Do not change the content of the first four rows of each worksheet (do not rename or remove the columns).
- Do not rename or move the worksheets in the template.
- Do not merge cells in the template worksheets.
- Do not use non-standard characters in the template; a Unicode character set must be used.
- Do not make any updates to the legend or valid values sheets.

#### 1.5 Reporting Template Worksheets

Each Reporting Template includes several worksheets which are described in Table 1.

An important concept regarding the organization of the Reporting Templates is that Station- and Sample-level information is captured in ***different worksheets*** than the Results-level information obtained from those samples.

- Example of Station-level information: coordinates
- Example of Sample-level information: sampling date, sampling depth
- Example of Result-level information: metal concentrations

[2] A list of ContractIDs can be found in Annex A.

**Table 1. Listing of the worksheets in the Reporting Templates.**

| Worksheet | Purpose |
|---|---|
| **GEO Reporting Template** | |
| Point Sample | Enter Station- and Sample-level information for samples collected at a point location (e.g., cores, grabs). |
| Towed Gear Sample | Enter Tow- and Sample-level information for samples collected a line location (e.g., trawl or dredge samples). |
| GeoResource_Results | Enter geological results for field collected samples. |
| Resource Information | Enter information about sampling locations that are specific to mineral-type. |
| Resource Assessment | Enter site-level information regarding resource assessment. |
| **ENV Reporting Template** | |
| Point Sample | Enter Station- and Sample-level information for samples collected at a point location (e.g., cores, grabs). |
| Towed Gear Sample | Enter Tow- and Sample-level information for samples collected a line location (e.g., trawl or dredge samples). |
| Chem_Results | Enter environmental chemistry results for field collected samples. |
| Biological_Results | Enter biological results. |
| **Worksheets in both GEO and ENV Reporting Templates** | |
| Legend | Field descriptions for each worksheet in the Template; identifies valid value fields, required fields, default values, maximum field lengths and default values for required fields. |
| Qualifiers | Stores definitions for qualifier codes included in GeoChem_Results. |
| ValidValues | Listing of allowable entries for fields constrained by valid values. |
| ValidValues_Analysis | Listing of allowable entries for the Category, Analysis, and Units fields in the GeoResource_Results and Chem_Results worksheets. |
| refQualifiers | Listing of example qualifier codes commonly used with analytical data. Examples are provided as a reference only. |
| StudyNotes | Enter any narrative information about the study in this worksheet. |
| ErrorChecker | The 'Error Checking' command executes a series of QAQC checks to ensure that the data set, as a whole, is reported correctly. |

#### 1.6 Legend worksheet

Strictly refer to the **Legend** worksheet of the Reporting Templates. Each of the fields in the Template are described in this worksheet and provides an important resource for understanding how to populate the Template. Table 2 provides descriptions of each of the columns included in the **Legend** worksheet.

**Table 2. Description of information provided in the Legend worksheets.**

| Column | Description |
|---|---|
| Worksheet | Indicates which Template worksheet the field is included in |
| Sample Worksheet | Indicates if the field is included in the 'Point Sample' worksheet, 'Towed Gear Sample' worksheet, or both. The purpose of the Sample Worksheet column is to avoid repetition of the fields that are common between the 'Point Sample' and 'Towed Gear Sample' worksheets. |
| Category | Indicates the Category of information; corresponds to the entries in Row 1 of the data entry worksheets. |
| Field | Name of the field included in the data entry worksheets |
| Data Type | Indicates the data type format in the DeepData database. |
| Definition | Description of the field. Includes important information for understanding the intended content of the field. |
| Valid Value Field | Indicates if the field is constrained to the list of valid values (included in the ValidValue and ValidValue_Analysis worksheets). See Section 8 for more information. |
| Required Field | Indicates if the field must be populated. Fields that are not required should be left null if true results were not obtained. Do NOT populate with placeholder values such as -8 or NR. |
| Maximum Length of Entry | Indicates the maximum number of characters allowed for the field. |
| Relevant Mineral Types | Indicates if the field is only relevant to certain mineral types. |
| Default values for Required Fields | For Required fields, indicates the default value to use if true results were not obtained. |

### 2. EXPLANATION OF KEY IDENTIFIER FIELDS

Fields that create the unique key for records in the **Point Sample** and **Towed Gear Sample** worksheet are:

**ContractID, Cruise Name, Leg Number, Station ID (or Tow ID), Sample ID**

#### 2.1 Expedition Identifiers

In the **Point Sample** and **Towed Gear Sample** worksheets, the key fields that identify data collected during an expedition are: **ContractID**, **Cruise Name** and **Leg Number**.

All data collected during an expedition (regardless of data type, sampling equipment or sampling method) are associated by assigning the same codes in these three fields.

#### 2.2 Station and Sample identifiers

The codes entered in the Station ID and Tow ID fields are identifiers for ***positions***, as indicated by different geographic coordinates.

Station ID is used in the **Point Sample** worksheet and is the identifier for samples collected from a point location (e.g., sediment core sample). Tow ID is used in the **Towed Gear Sample** worksheet and is the identifier for samples collected from line locations (e.g., tows or trawls).

The codes entered in the Sample ID field are identifiers for samples collected (e.g., depth intervals from a sediment drill sample).

In this context, a Station ID does NOT identify a feature (e.g., a specific chimney edifice). If a group of Station IDs are associated with a feature, this should be entered in the Station Description field. In Annex B, Example 1, the same feature (Edifice 123) was sampled on 3 different days; the coordinates for each Station ID are slightly different, and each different ROV dive is given a different Station ID; an entry is provided in the Station Description field to allow grouping by the sampled feature.

Conversely, Example 2 is a drill sample that was sub-sampled into intervals. The Station ID, Sample Date and coordinates are the same, and the 4 different depth intervals are indicated in the Sample ID and Upper/Lower Depth fields.

For a single expedition, the combination of Station ID+Sample ID (or Tow ID+Sample ID) fields must be coded to identify unique field-collected samples.

Using the same Station ID (or Tow ID) associates all data collected at that location; using different Sample IDs differentiates samples collected that are different matrices (sediment or water or rock), collected on different dates and/or different depths.

**See Section 6 for special instructions for coding Station and Sample identifiers for CTD casts.**

#### 2.3 Relating Information in Different Worksheets

For GEO Template:

- Station ID+Sample ID entries in the **Point Sample** worksheet must match to entries in the **GeoResource_Results** worksheet.
- Tow ID+Sample ID entered in the **Towed Gear Sample** worksheet must match to entries in the **GeoResource_Results** worksheet.

- Station ID or Tow ID entries in the **Resource Information** worksheet must match to entries in the **Point Sample** or **Towed Gear Sample** worksheet.
- Exploration Block ID or Exploration Subarea entries[3] in the **Resource Assessment** worksheet must match to entries in the **Point Sample** or **Towed Gear Sample** worksheet.

For ENV Template:

- Station ID+Sample ID entries in the **Point Sample** worksheet must match entries in either the **Chem_Results** worksheet or the **Biological_Results** worksheet.
- Tow ID+Sample ID entries in the **Towed Gear Sample** worksheet must match entries in either the **Chem_Results** worksheet or the **Biological_Results** worksheet.

The Error Checker will ensure that Station ID/Sample ID/Tow ID in the sample-level worksheets are represented in the results-level worksheets [an exception to this is tow samples that only have video(s) associated with them].

#### 3. INSTRUCTIONS FOR ASSIGNING STATION IDS

Different Station IDs should be assigned when the Actual Latitude/Actual Longitude are different. The exception is CTD casts (see Section 6 below).

Preferably, the Actual Latitude/Actual Longitude should be measured with the instrument at the seafloor (bottom samples), or the sample collection point (for mid-water sampling). The location of the instrument recording coordinates must be specified in the Position of GPS/Transducer field.

The formats for coordinates must be provided in Decimal Degrees (DD); all other coordinate formats are not acceptable [e.g., Degrees/Minutes/Seconds (DMS)].

Information on station locations must be made to the most accurate geographical and depth measurements, whenever possible.

#### 4. INSTRUCTIONS FOR ASSIGNING SAMPLE IDS

When samples are collected at different depths, each sample interval should be assigned a different Sample ID. See Annex B, Example 2.

[3] A shapefile with the Exploration Block ID and Exploration Subarea entries can be accessed from the 'Downloads' tab of the DeepData website (https://data.isa.org.jm/isa/map/).

Different matrices sampled must have unique Station IDs+Sample IDs (e.g., a sediment and porewater sample collected at the same station should be considered two separate samples, and Sample IDs such as 01S and 01W should be assigned). See Annex B, Example 3.

Similarly, if total and dissolved fractions are measured in the same field-collected water sample, each fraction must be assigned a different Sample ID (e.g., 01T for the total fraction and 01D for the dissolved fraction). See Annex B, Example 4.

Sample IDs for geochemistry samples must be different from the Sample IDs given to biological samples. See Annex B, Example 5.

Since **Point Samples** and **Towed Gear Samples** are captured in different worksheets, it is important that the Station IDs and Sample IDs do not overlap between the two worksheets. So, if both Point Samples and Towed Gear Samples are collected during the same Cruise, the same Sample ID must not be used in both the Point Sample worksheet and the Towed Gear Sample worksheet.

#### 5. CODING SAMPLES WITH BOTH CONFIDENTIAL AND PUBLIC DATA

Most cruises will collect samples that will be designated as EITHER confidential OR public data. These **separate** samples will be differentiated by being included in EITHER the GEO Template OR the ENV Template, respectively.

There is the special example of **individual** samples that include both confidential AND public data. An example is a box core sample that is sub-sampled for nodules and sediment and both are analyzed. In this case, the nodule results must be entered in the GEO Template; the sediment results must be entered in the ENV Template. To identify the samples as sub-samples:

- In both templates, enter the word "Paired" in the Sample Collection Method field.
- For these 'Paired' samples, use the same ContractID/Cruise Name/Leg Number/Station ID in both templates.
- Use the same root Sample ID in both templates and add a "P" suffix (for paired) to the Sample ID in the ENV Template.

#### 6. INSTRUCTIONS FOR ENTERING CTD CAST DATA

CTD data is publicly accessible and therefore must be entered in the ENV Template.

Samples and sensor readings from the same cast should have the same Station ID, ***even if there are unique coordinates for each Sample ID*** (this is an exception to the rule that each set of unique coordinates should have a unique Station ID; see Section 3).

For water samples collected from Niskin bottles, collected at the same depth as a sensor reading, use the same root Sample ID, and add a "S" suffix (for sample).

In the **Chem_Results** worksheet, results from CTD sensor readings must have Category = 'Water Properties'. The depth and sample date for CTD sensor readings are entered in the CTD_Depth and CTD_SampleDate fields.

In the **Point Sample** worksheet, samples associated with CTD casts must have Sampling Device = 'CTD sensor' and Profile = 'Cast'.

See the Template Example Package for specifics on entering CTD cast data in the ENV Reporting Template.

### 7. QUALIFIER CODES

Analytical data is accompanied by qualifier codes that flag results with certain problems and scenarios (e.g., U = Analyte was not detected at or above the reporting limit; J = estimated result). The laboratory will assign qualifier codes to individual results to assist users with interpreting the data. There may also be an independent 3rd party validation (data quality assessment) in which the laboratory data are scrutinized, and the validators add validation qualifier codes.

Examples of qualifier codes commonly used with analytical data are included in the **refQualifiers** worksheet. These example codes are provided for reference only.

The qualifier codes accompanying the results from the analytical laboratory should be captured in the 'Qualifier' column in the **GeoResource_Results** (GEO Template) or **Chem_Results** (ENV Template) worksheet.

Qualifier codes in the **GeoResource_Results** or **Chem_Results** worksheets must be included in the **Qualifiers** worksheet and defined.

### 8. VALID VALUE LISTS

The purpose of the Valid Values lists is to standardize and constrain the entries in a particular field so the data can be more consistently filtered, sorted, and analyzed.

Fields constrained by valid values are identified in the **Legend** worksheet, and in row 2 or 3 of the **Point Sample**, **Towed Gear Sample**, **GeoResource_Results/Chem_Results** and **Biological_Results** worksheets.

#### 8.1 ValidValue_Analysis Worksheet

- Entries in the 'Category' and 'Analysis' columns of the **GeoResource_Results/Chem_Results** worksheet must exactly match the entries in the **ValidValues_Analysis** worksheet.
- Entries that are not included in this list will be identified by the Error Checker (see Section 10) and an error message will be given so that the invalid entry can be easily identified and corrected.
- Entries in the 'Units' column of the **GeoResource_Results/Chem_Results** worksheet must match one of the options in the UnitsSolid and UnitsWater fields in the **ValidValues_Analysis** worksheet.
- The **ValidValues_Analysis** worksheet includes the current list of analytes for the ISA DeepData database, however, not all analytes must be reported for each template and matrix type.
- New items may be added to the **ValidValues_Analysis** worksheet, but existing entries must not be modified. New items must include information in the Description field. Before adding an item to **ValidValues_Analysis**, be sure to do a thorough search to see if the item is already on the list (i.e., use the Find command; search for the item using different key phrases). New items in **ValidValues_Analysis** will be reviewed by the ISA Data Management Team and incorporated into the master list, if approved.

#### 8.2 ValidValue Worksheet

- The allowable entries for all other valid value fields (i.e., not including Category, Analysis, and Units fields from the **GeoResource_Results/Chem_Results** worksheet; see Section 8.1) are included in the **ValidValues** worksheet.
- Entries in these valid value fields must exactly match the entries in the **ValidValues** worksheet.
- Entries that are not included in this list will be identified by the Error Checker (see Section 10) and an error message will be given so that the invalid entry can be easily identified and corrected.
- The ValidValue list includes special codes if information for valid value fields is not available (e.g., 'Not Applicable' or 'Not Reported'). The Error Checker will only accept these values as correct entries.
- It is anticipated that the current valid value list for 'Sampling Device' is not comprehensive, therefore it is acceptable to enter a Sampling Device that is not included in the valid value list – in this case ignore the Error Checker error. New items will be reviewed by the ISA Data Management Team and incorporated into the master list, if approved.

### 9. INSTRUCTIONS ON LINKING RESULTS TO IMAGE FILES

Image files are an important aspect of documenting and characterizing the deep seabed environments. Examples of image files: video tracks taken from an ROV; photographs of organisms retrieved from collected samples, a frame from a video displaying a feature or organism observed in the water column.

Information on individual photos and videos is required to allow linkages to specific samples, tows, or results (i.e., organisms), which can then be accessed via the DeepData website Photo/Video Gallery tab.

Linkages between samples/results and image files must be made in BOTH the Reporting Template and the associated Metadata Template.

In the Reporting Template, information on photo/video files is entered in the following worksheets and fields:

**Point Sample** - Photo File Name

**Towed Gear Sample** - Video File Name

**Biological_Results** - Photo File Name, Video Frame Code, Video photo frame file name

If multiple image files are submitted for a single sample or result, include multiple entries in these fields, separated by commas.

When information on photos/videos is entered in the Reporting Template, they must also be logged into the **PhotoVideoFiles** worksheet of the associated Metadata Template.

The Metadata Template:

- allows storage of additional information on the photo/video files (e.g., Camera Specs, DateTime Stamp, description) and,
- provides the mechanism for linking and accessing photo/video files through the website Photo/Video Gallery interface.

See Section 13.5 for specific instructions on populating the Metadata Template with information on image files.

### 10. ERRORCHECKER

The ‘Error Checking’ command, in the **ErrorChecker** worksheet executes a series of QAQC checks to ensure that the data set, as a whole, is reported correctly.

A Checker Report is generated with messages describing the errors found, so the user can easily identify problems. Note that three different types of error messages are generated: Pass, Fail, Alert.

After resolving errors or making any modifications, it is necessary to re-run the ‘Error Checking’ command (this is to ensure that the Checker Report is updated to reflect the most recent changes in the Templates).

The Template may not be accepted by the ISA if the Checker Report includes any Fail error messages.

### Reporting Templates – Specific Instructions

#### 11. Specific Instructions for the GEO Reporting Template

This section provides specific instructions for entering confidential data into the GEO Reporting Template. The general instructions for entering sampling information (i.e., Point Sample and Towed Gear Sample) apply to the GEO Reporting Template, and this section focusses specifically on entering resource assessment results that are specific to the GEO Reporting Template.

##### 11.1 Data Entry Worksheets

When entering results from geological prospecting and sampling into the GEO Reporting Template, Contractors provide information in five different worksheets:

1. **Point Sample** - Cruise, Station and Sample information for samples collected at point locations (e.g., drill samples)
2. **Towed Gear Sample** - Cruise, Towed Gear location, and Sample information for samples collected at tow locations (e.g., dredge samples)
3. **GeoResources_Results**: Results from geochemical analysis are captured in this worksheet - Station ID (or Tow ID), Sample ID, Category, Analysis, Result, Units, Qualifier, Limits of Detection (LOD), Analytical Technique, Lab Matrix, Total or Dissolved, Measurement Basis, Laboratory, Remarks.
4. **Resource Information** – information associated with specific mineral types to characterize resources at sampling locations (e.g., Hydrothermal Activity, Nodule abundance, Crust Thickness)
5. **Resource Assessment** – information associated with resource assessment for specific mineral types, at a site-level (e.g., Extensions of PMS site, Inferred Tonnages, Area of Crust Coverage)

##### 11.2 Data and Information to Submit for Geological and Resource Assessment

Mineral exploration aims at the identification and assessment of mineral resources. Identification of mineralized areas and subsequent definition of deposits with potentially economic ore grades, and earliest spatial reduction of the exploration (i.e. license) area for cost reduction are of key interest of land-based exploration, and the same approach is required for economic seafloor exploration. All three mineral resources in the Area have specific characteristics for their formation and enrichment to deposit-sizes that largely depend on the regional to local geological and oceanographic conditions. Geological models were successfully developed which reliably describe the conditions of formation for all three types. However, ore deposit formation requires specific processes and favorable conditions, and their detailed documentation is critical for deposit characterization.

Specific local characteristics and mineral endowment as well as overall variation of geological and oceanographic conditions in un- or less-mineralized regions of the Area can ideally be described by the following (categorized as Unstructured or Structured data):

**Unstructured**

1. Detailed swath bathymetric survey(s) of the license area with resolutions better than 20 meters for areal coverage of manganese nodules (PMN) and better than 5 meters for local mineral enrichments (polymetallic sulphides, PMS, cobalt-rich crusts, CFC). Detailed and high-precision swath bathymetry allows for spatially high-resolution sample localization and characterization.
2. Detailed geophysical surveys for subsurface identification and modeling: Gravimetry, magnetics, (backscatter) sonar (PMN, PMS, CFC), electromagnetics (PMS, CFC), seismics (3D, PMS).
3. Detailed geological mapping and photo documentation of the identified deposit (towed video sled, ROV, AUV, HOV).

These unstructured data types are submitted as raw files and cataloged in the Metadata Template (see Section 13).

**Structured**

1. Resource assessment and classification by determination of coverage density (PMN) and by drilling (PMS, CFC).

These structured data types are submitted in the GEO Reporting Template format.

Analysis and assessment of these data ultimately will result in:

- Scientific assessment of geological exploration data and deposit modeling, and,

- Detailed geological characterization and assessment, and definition of prospective areas according to geological evolution of the license area: Structural geology, volcanism, sedimentation processes, mineral precipitation, remobilization and redistribution.

Implementing these data collection efforts during exploration facilitates the assessment of resources, deposit characterization and classification. Survey data are submitted in raw format and logged into the Metadata Template (e.g., bathymetric surveys); analytical data are re-formatted to DeepData's confidential GEO Template to be incorporated to the DeepData database as structured data. The documentation assists in the compilation, understanding, interpretation and refinery of exploration data to official mineral title application forms, market analysis of mineral resources and required technical resource classification reports (e.g., NI 43-101, CRIRSCO, ISA Reporting Standard template).

#### 11.3 Provisional Results

Submissions to DeepData should include provisional results during exploration. It is recognized that the assessment, characterization, and deposit definition are usually a long and iterative process. Provisional results should be submitted to DeepData, with the understanding that the different identified sites and areas will be revisited continuously, geological and analytical results may differ over short distances and the assessment will be refined and re-submitted as exploration and sampling continues, and analyses are refined to a greater level of detail. Information on the status of characterization and the deposit identification process are indicated by the Cruise and Station Information, the Sample Date; specific indications of the status of analytical results can be indicated in the Qualifier field of the **GeoResource_Results** worksheet. It is understood that entries in certain fields will be incomplete when the status of knowledge is interim.

When results from a previously submitted Reporting Template are revised (e.g., Qualifiers are added; additional parameters are added; Results are changed), the Reporting Template must be re-submitted. **Re-submissions must include the entire dataset, not just the records that have been updated.** See Section 14.1 for the detailed procedures for submitted revised Reporting Templates.

Note that previous submissions will be removed from DeepData and archived; the re-submitted dataset will replace the original version. A record of the progression of deposit definition and resource assessment over time will be stored in the DeepData Archive.

#### 11.4 Data Requirements

The **ValidValues_Analysis** worksheet lists information on a vast variety of different chemical, physical and compositional analytical categories. These include element analyses, geotechnical parameters, mineralogical compositions, physical properties, and nutrient compositions. The type of samples collected during seafloor exploration is quite variable and includes organic and inorganic materials, solid and diverse volcanic rock samples, soft and indurated sediment samples of different origin (e.g., volcaniclastic, terrigenous, biogenic), highly diverse mineral precipitates; pore water, bottom-interface water and open ocean seawater, and also mixtures and intergrowths of all the former. It is evident that not all the analytical categories are applicable to every single sample, that a number of samples need severe sample processing for a proper analysis of single categories, and that the sampling process as well as the sample treatment can severely alter the sample composition.

This version of the GEO Template does not specify the required measurements over an exploration period, nor does the Error Checker verify that the required measurements have been submitted. This verification will be conducted on the data compiled for the entire exploration period (i.e., the checks will be conducted by the ISA Data Management Team on the data compiled in the database for the contract).

#### 11.5 Data Reporting Specifications

An increasing number of chemical and physical parameters can be analyzed in the field, and both on-line measurements and lab measurements (e.g., pH values and element concentrations, respectively) need to be differentiated by providing information in the Sample Collection Method and Sampling Device fields. It is necessary to supply information on the sampling methods and analytical techniques applied and to report critical detection limits as well as analytical uncertainties, in the LOD (Limit of

Detection) and Qualifier fields, respectively. Analytical caveats, however, should not prevent from reporting as continuing exploration helps to review, refine and further state the analytical information more precisely. Also, the potential general variability in compositions (at least for some geological samples) will provide important information on natural variability within the license area, with consequences for the exploration as well as for environmental baseline surveys.

The detection of physical and geotechnical parameters will largely influence any potential development of mining technologies and future mining strategies. Measurements are critical for successful technical solutions and improve the development of low-impact collector, mining and recovery systems. Proper selection of the type of analysis, its execution, critical operation analysis and documentation of the analysis and results are necessary and therefore suggested.

For each Category/Analysis reported in the **GeoResources_Results** worksheet, it is required to report in the following fields: Category, Analysis, Result, Units. It is advantageous to also provide information in the following fields: Analytical Technique, Qualifier and LOD (Limit of Detection). In addition, it is important to provide additional information for the following Categories, to aid in data interpretation:

- For the Elements and Rare Earth Elements Category - Loss on Ignition (wt%) should be included
- For the Nodule Properties Category
    - Analyzed Segment should be reported, to indicate whether the nucleus, interior or rim was the analyzed segment.
    - Layer Sampled (in the **Point Sample** worksheet) should be reported, to indicate the layer sampled, relative to the surface of the nodule, in mm).
- For the Mineralogy Category
    - It is recognized the quantitative minerology is a complicated process and is not necessary to be determined for every sample. Percentages of specific minerals (e.g., 'Minerals – Feldspar' need only be reported when available and should be accompanied with notes on analytical conditions [enter in the Remarks field; for example: timing, x-ray conditions (shaking or light), method preparation].

At this time, laboratory QA/QC results (e.g., method blanks, lab replicates) should not be incorporated in the Reporting Template. This information can be submitted as raw data via the Metadata Template (see Section 13).

Analytical results stored in the **GeoResources_Results** worksheet must be made to the highest precision and qualified confidence level, whenever possible. Analytical results must be reported in standardized units, which are listed in the **ValidValues_Analysis** worksheet.

Other specifics to note regarding the Matrix field:

- Sediment porewater is interstitial water, which is water extracted from the sediment column, NOT water collected at the sediment/water interface.
- For sediment porewater samples, Upper Depth and Lower Depth entries will reflect the sediment sampling depth (although Matrix Type = porewater).

- All analytical results for solid matrices (i.e., rock, sediment) must be reported on a dry weight measurement basis.
- In the GEO Reporting Template, the Matrix Type must be assigned as either:
  - 'Mineable Substrate – Nodules'
  - 'Mineable Substrate – Crust'
  - 'Mineable Substrate – Sulphide Mounds'

### 12. Specific Instructions for the ENV Reporting Template

This section provides specific instructions for entering publicly available data into the ENV Reporting Template. The general instructions for entering sampling information (i.e., Point Sample and Towed Gear Sample) apply to the ENV Reporting Template, and this section focusses specifically on entering biological results that are specific to the ENV Reporting Template.

For Station ID + Sample ID that have associated results in the **Biological_Results** worksheet, the Matrix Type must be assigned as 'Biological sample'.

#### 12.1 General

For entering results from biological sampling into the **Biological_Results** worksheet of the ENV Reporting Template, the following six categories of information are available:

1. **Taxa ID** (i.e., the taxonomic identification of individual organisms; Class, Family, Species, etc.; identification method and status)
   a. The entries in the Species field should be the species name without the author citation (e.g. "hoffmeisteri" not "hoffmeisteri Claparède, 1862"). The author citation should be entered in the 'Taxonomic author citation' field.
   b. Note that the 'Scientific Name' field is a Required field and is a key field for verifying taxonomic identifications against the World Register of Marine Species (WoRMS) taxonomic database (see below).
2. **Taxonomy Information** (i.e., information regarding traceable data or evidence from which the taxonomic identification was obtained, e.g., voucher specimens, animal tissues, and DNA sequence information)
3. **Identifier Information** (name, email, institution, whether the identifier is a taxonomist)
4. **Ecology** (e.g., number of individuals collected, biomass, etc.)
5. **Organism Details** (e.g., sex, life stage of individual organisms, etc.)
6. **Photo/Video Information** (if applicable, this information is captured to allow a linkage to the file collection Library, enabling database users to quickly identify and access photos/videos of a specific sample or organism using the DeepData website).

#### 12.2 Details on Identification Methods and Taxonomy Information

Following the guidance provided by the Legal and Technical Commission (ISBA/25/LTC/6/Rev.1, Annex I, para. 47), more accurate and consistent taxa identification is provided by two sets of information: morphological descriptions (e.g., images, drawings) and genetic sequences. For new species descriptions, the two sets of information are then published in scientific journals as well as added to international databanks and curated collection facilities (voucher specimens, molecular samples, including DNA extracts).

There are three different types of Identification Methods that can be entered in the Reporting Templates: **Morphological ID**, **Genetic ID** and **Remote ID**. Specific information is required to be reported for each Identification Method (see details below). In addition, there is an option to indicate 'combination' Identification Methods (see the **ValidValue** worksheet); this allows data providers to indicate instances where multiple identification methods were used (i.e., genetic + morphological; Remote ID + genetic/morphological).

Under the Taxonomy Information category, the data provider is encouraged to report on taxonomic databases [e.g., World Register of Marine Species (WoRMS), GenBank, etc.] accessed to record the information that would allow an end user to verify the scientific name reported; this information is entered in the following fields: *Taxonomic Database*, *Database Taxa ID*, *LSID*, *INSD accession number*, as well as details on vouchers (see specific details below). **The purpose of tracking the ancillary data associated with a taxonomic identification is to allow regulators (and other data users) to:**

1) **locate specimens;**
2) **track information on how identifications were determined; and**
3) **validate the identification.**

It is critical to correctly populate the taxonomic categories (i.e., Kingdom, Phylum, etc), with respective scientific name under each taxonomic category, up to the lowest taxonomic rank that can be determined.

##### 12.2.1 Morphological ID

If identification was made by an expert using morphology and phenotypic characteristics (Identification Method = Morphological ID), the following Template fields should be populated:

a. **Taxonomic Database** – Reference database used to verify the reported taxonomic name (e.g., WoRMS). For a listing of available databases, refer to Table 1 in: Rabone *et al.* 2019.
b. **Database Taxa ID** - The associated taxon identifier from the taxonomic database (e.g., if using the WoRMS website, the AphiaID code would be entered; note that a Database Taxa ID is required for species level as well as higher-level taxa identification),
c. **Taxonomic Status** – Status of taxon details, as indicated in the taxonomic database; in case of pending/interim identifications, "NA" should be entered).
d. **Taxonomic Identification Qualifier** - A controlled value to express the determiner's doubts about the Identification (see **Legend** worksheet, Templates Example Package, and Horton *et al.* 2021).

e. **Notes on Taxonomic Identifications –** Additional remarks on identification status, explaining taxonomic identification qualifier (see Templates Example Package for example entries).
f. **LSID** – the Life Science Identifier

The Life Science Identifier (LSID) is critical, as it provides a 'naming standard underpinning for wide-area science and interoperability' (LSID, n.d.). Data providers are encouraged to use the WoRMS database to verify taxonomic nomenclature because WoRMS tracks changes to taxonomic names, and the ISA will use WoRMS to verify the taxonomic information submitted by Contractors. The WoRMS website provides a Taxon Match tool, to automatically match a species or taxon list with the WoRMS database. Guidance for using this tool can be found at:

1) http://www.marinespecies.org/aphia.php?p=match
2) https://www.youtube.com/watch?v=Mk1IiEwvsF8 (from 5min48sec to 13min55sec)

##### 12.2.2 Genetic ID

If identification was made using genetic methods, such as DNA barcoding, (Identification Method = Genetic ID), the following Template fields should be recorded, for a Final identification:

a. **Taxonomic Database** - Database used for data submission (e.g., DDBJ, ENA, NCBI)
b. **INSD accession number** - INSD accession number(s) assigned by the data submission process of the DDBJ, ENA or NCBI databases.
c. **Description of DNA Marker** - describe the gene sequence (e.g., COI, 16S genes, 18S genes, etc.)
d. **Tissue Descriptor –** information on the tissue type submitted for DNA analysis.
e. **LSID** – the Life Science Identifier (see above)

There are three members of the INSD (the International Nucleotide Sequence Databases): DDBJ, ENA, NCBI (=Genbank). Only one accession needs to be made at any of the three databases and a single accession number is propagated to all three sources. Further information about DNA accessions is available on the DDBJ (DNA Data Bank of Japan) website: http://www.ddbj.nig.ac.jp/sub/acc_def-e.html.

In cases where the identification is made solely by DNA sequencing, information on phylogeny should be provided to the lowest possible taxonomic rank (i.e., entries in the high-level taxonomic categories; Kingdom, Phylum, etc.), as well as "Morphotype", if the Identification Status is Interim or Final at the time of Template submittal. The WoRMS LSID should be entered for the lowest taxonomic level determined.

##### 12.2.3 Remote ID

If identification is made remotely, i.e. using video or image observations, without physical specimens (Identification Method = Remote ID), the following Template fields should be recorded:

a. **PhotoVideo Information –** Images that are associated with the organism identified by Remote ID should be submitted to DeepData and associated with the specific organism using the following fields: Photo file name, Video frame code, Video photo frame file name.[4][5] Note that multiple images can be submitted for a single organism identification using comma separated entries (see section 9 on instructions for linking results to image files).

b. **Taxonomic Database** – If the images used in the remote identification were submitted to an image database, include the database in the Taxonomic Database field (e.g., Biigle).

f. **Morphotype** - An informal group of taxa with similar or identical morphology; often preferred to describe morphological variation within a species, especially if a formal description as a variety appears not strongly supported or premature. *For cross-referencing purposes, the identifier entered in the Template must match exactly the morphotype title/identifier in the Taxonomic Database referenced.*

g. **LSID** – the Life Science Identifier (see above)

#### 12.3 Details on Vouchers

Following the guidance provided by the Legal and Technical Commission (ISBA/25/LTC/6/Rev.1, IV.A.19; IV.B.22), "*Specimens **must** be archived for comparison with taxonomic identifications from other sites and to understand the details of changes in the composition of species over time. If species composition does change, it might be subtle, and reference back to the original animals (where there might only have been a putative identification) is essential. It is recommended that samples be archived as part of national or international collections and that appropriate funding be identified to facilitate this*."

A specimen voucher is the verifiable reference that can be used (and revisited) to confirm an identification. This means that the identifications can be compared between surveys, by checking the voucher specimens. There are two types of voucher specimens, as follows:

(1) Type specimens – upon which formally accepted names of taxonomic units are based;
(2) Taxonomic support specimens – which document identifications in taxon-based studies other than nomenclatural studies.

DeepData requires a taxonomic support specimen to be retained for each identification made. One or two representative organisms must be vouchered, given a code, and stored. **It is suggested that the samples are submitted to a collection registered on this site:** https://www.gbif.org/grscicoll

The following information about the voucher must be captured in the Template[6]:

[4] See Section 9 and 13.5 of this guidance, Section 3.2.4 of the User Manual, and the Template Example Package for detailed instructions on submitting photo and video files and recording information in the Metadata Template to document these files.
[5] Note that procedures for submitting image files is currently under review and revisions are anticipated in the next update of the Reporting Templates.
[6] If multiple vouchers are submitted, include multiple entries in these fields separated by commas.

a. **Voucher status –** a description of the stage of submission and approval/acceptance of the voucher specimen (e.g., submitted, frozen, rejected, destroyed, accepted, pending approval);
b. **Voucher code -** Code assigned by the submission process the voucher is submitted to; identifies the stored voucher specimen;
c. **Voucher institution code –** Code for the museum/institution where voucher specimen is stored.

#### 12.4 Submitting Ancillary Information

Data providers are strongly encouraged to submit the following types of information that are associated with taxonomic identifications:

1) Images of organisms obtained during field sampling, and/or laboratory studies (e.g., microscope photos, drawings);
2) Publications;
3) Protocols used during sampling and identification, including adjustments to the Polymerase Chain Reaction (PCR) protocols.

Files containing this information are submitted as unstructured data, using the Metadata Template to log and categorize files. See Section 13 of this guidance, Section 3.2 of the User Manual, and the Templates Example Package for detailed instructions on submitting ancillary information.

#### 12.5 Provisional Results

It is recognized that identifying species can be a long and iterative process. Submissions to DeepData accommodate provisional taxonomic results, with the understanding that the results may be refined and re-submitted at a later date. Information on the current status of the identification process should be included in the following Template Fields: *Taxonomic Identification Qualifier*, *Identification Date*, *Identification Status* (i.e., Final or Interim) and *Notes on Taxonomic Identifications*; for information on the use of *Taxonomic Identification Qualifiers* (e.g., *stet.*, *indet.*, *inc.*, *aff.* and *cf.*) see examples and explanations provided in the **Legend** worksheet, Templates Example Package, and Horton *et al.* 2021. It is understood that entries in certain fields will be incomplete when the Identification Status is Interim (e.g., entry in the 'Database Taxa ID' should be set to 'Not Available').

When results from a previously submitted Template are revised (i.e., Identification Status and Identification Date is changed; Taxa ID information is updated; **Chem_Results** are revised), the Template must be re-submitted. **Re-submissions must include the entire dataset, not just the records that have been updated.** See Section 14.1 for the detailed procedures for submitting revised Reporting Templates. Note that previous submissions will be removed from DeepData and archived; the re-submitted dataset will replace the original version. A record of the progression of an identification over time will be stored in the DeepData Archive.

#### 12.6 Ecological and Organism details

Information on taxa abundance (i.e., number of individuals) and biomass is entered under the Ecology section of the Biological results. Other calculated ecological indices (e.g., Relative abundance, Relative Dominance, Taxon Density) may also be provided, but are not required to be submitted by Contractors in the Template. If this information is reported in the Template, Contractors are encouraged to provide details on the calculation procedures used, either in the 'Additional notes about ecology' field, the StudyNote tab, or in associated documentation submitted as unstructured data. Other biological indices (e.g., spatial extent, density of particular taxa, diversity) that have been determined should also be submitted as unstructured data. When available, organism details (e.g., life stage, sex, etc.) should be entered in the template for each taxa.

#### 12.7 Chem_Results in the ENV Template

If there is ancillary data collected along with biological sampling (e.g., grain size information from a sediment sample, presence of coral assemblages), this information should be entered in the **Chem_Results** worksheet. Specifics to note about entering results in the **Chem_Results** worksheet:

- For the CFC mineral type, there is a requirement to measure parameters in water samples collected as close as possible to the rock/water interface (epibenthic boundary layer). For these samples, the Habitat Type must be assigned as 'Epibenthic boundary'.

- For the PMN mineral type, there is a requirement to measure parameters in the pore water matrix. For these samples, the Matrix Type should be assigned as 'Sediment Porewater filtered' or 'Sediment Porewater unfiltered'.
    - For these samples, Upper Depth and Lower Depth entries will reflect the sediment sampling depth;
    - Sediment porewater is interstitial water, which is water extracted from the sediment column, NOT water collected at the sediment/water interface.
- All analytical results for solid matrices (i.e., rock, sediment) must be reported on a dry weight measurement basis.

### METADATA TEMPLATE & DATA SUBMISSION

#### 13. METADATA TEMPLATE

The Metadata Template is used when submitting all data (structured and unstructured) to the International Seabed Authority. The purpose of this template is to categorize and document information about the files submitted over the website, and to streamline the receipt, import, and classification (confidential, non-confidential) of both structured and unstructured data provided by the Contractors.

In addition to this guidance, the User Manual (Section 3.2) and the Template Example Package should be referenced for instructions on populating the Metadata Template.

##### 13.1 File Naming Convention for Metadata Template

Use the following naming convention for the Metadata Template file:
MetadataTemplate_ + ContractID + Contractual Year (e.g., MetadataTemplate_DORDPMN12015).

The file name may not exceed 120 characters.

##### 13.2 Modifying the Template Structure

The Template structure is designed to be compatible with ETL procedures to append results into the DeepData database. As such, the structure is not flexible:

- All fields in these worksheets are text and should remain as text.
- Do not change the content of the first three to four rows of each worksheet (do not rename or remove the columns).
- Do not rename or move the worksheets in the template.
- Do not merge cells in the template worksheets.
- Do not use non-standard characters in the template; a Unicode character set must be used.
- Do not make any updates to the legend or valid values sheets.

##### 13.3 Metadata Template Management

The Contractor fills out a **single** Metadata Template for all data associated with the Contractual Year (i.e., all data associated with an Annual Report).

The same Metadata Template file is submitted with multiple submittals (i.e., update a single Metadata Template file for all data associated with an Annual Report). Include the current version of the Metadata Template with each compressed (zip) file submitted (i.e., EACH zip file must include a Metadata Template file).

Each compressed (zip) file logged in the Metadata Template must contain files associated with ONLY a single theme (see **ValidValues** worksheet for a listing of themes).

Reporting Templates must be submitted with only one Reporting Template per zip file.

Within a single Metadata Template file, there cannot be two 'Zip File Name' entries OR individual file names that are the same.

#### 13.4 Instructions for Populating

Required fields must be populated. Other fields can have null entries.

A description of the worksheets included in the Metadata Template file is shown in Table 3.

**Table 3. Description of worksheets in Metadata Template.**

| Worksheet Name | Descriptions |
|---|---|
| **Legend** | Field descriptions for each worksheet in the Template |
| **Annual Report** | Complete this worksheet when submitting Annual Report document(s) in a compressed (zip) file. |
| **ReportingTemplates** | Complete this worksheet when submitting Reporting Template files in a compressed (zip) file. |
| **NonTemplateData** | Complete this worksheet when submitting all other data files (i.e., data that is not accommodated by the Reporting Templates), including compressed files containing photos and videos. |
| **PhotoVideoFiles** | If the NonTemplateData worksheet includes files with Theme = Photo_Video, then complete this worksheet with information on the individual photo and video files. |
| **ValidValues** | Listing of allowable entries for fields constrained by valid values. |
| **ErrorChecker** | The 'Error Checking' command executes a series of QAQC checks to ensure that the data set, as a whole, is reported correctly. |

The individual files included in the compressed (zip) files do not need to be cataloged in the Metadata Template, **except for files submitted under the Photo_Video Theme.**

#### 13.5 Photo_Video Theme

Information on individual photos and videos is required to allow linkages to specific samples, tows, or results (i.e., organisms), which can then be accessed via the DeepData website Photo/Video Gallery tab.

Linkages between samples/results and image files must be made in BOTH the Reporting Template and the associated Metadata Template. See Section 9 for specific instructions for populating the Reporting Template with information on image files.

Six different scenarios have been identified where photos or videos may be procured and accommodated in the Metadata Template, Reporting Templates, and the geodatabase structure. The six photo/video scenarios identified in Annex C provide examples of appropriate data entry in the Metadata Template for photos and videos.

Multiple image files are included in a compressed file (.zip), and this file is logged in the **NonTemplateData** worksheet with an appropriately named 'Zip File Name' (see Section 13.6), and Theme = Photo_Video. The ***individual*** image files included in the zip file are cataloged in the **PhotoVideoFiles** worksheet, including the name of the individual file in the 'VP File Name' field (e.g., DSCN2095.jpg). Also enter information in the Station ID, Tow ID Sample ID, OrgNum and Image Number fields to link individual photos and videos to specific point samples, tow samples, or results. Each 'VP File Name' is associated with a 'Zip File Name', that was logged into the **NonTemplateData** worksheet.

If multiple image files are submitted for a single sample or result, differentiate these by entering an arbitrary identifier (e.g., 1, 2, 3) in the Image Number field.

If photos/videos are not associated with specific samples or results, then enter 'NA' in the Associated Results field and leave the Station ID, Tow ID, Sample ID, OrgNum and Image Number fields null.

Records in this **PhotoVideoFiles** worksheet must be unique based on the following fields:
Station ID + Sample ID + Tow ID + OrgNum + VP File Name + Image Number

#### 13.6 File Naming Convention for Data Submissions

A data submission is a single zip file. Each zip file includes a Metadata Template file and one or more data files. The name of the data submission file is entered in the Zip File Name field in the appropriate worksheet in the Metadata Template. The zip files must adhere to a naming convention[7].

Each **'Zip File Name'** within a Metadata Template for a Contractual Year must be unique.

The file name may not exceed 120 characters.

Use the following naming convention for Reporting Template submissions:

ContractID + Contractual Year_ + master file name_ + descriptive details
For example: BGRPMN12015_Geo_Template_2014rock.zip

Use the following naming convention for other data submissions:

ContractID + Contractual Year_ + Theme_ + descriptive details
For example:

[7] The file names are important as they are referenced in automated data processing procedures, which are used to accurately log and categorize the various types of data types submitted.

- DORDPMN12015_AnnualReport_appendices.zip
- JOGMECCRFC12016_Bathymetry_CruiseA.zip

**Submitted files must not have the same file name as previously submitted files.**

##### 13.7 ErrorChecker

The 'Error Checking' command executes a series of overall checks to ensure that the data set, as a whole, is reported correctly.

A Checker Report is generated so the user can easily identify problems. Note that three different types of error messages are generated: Pass, Fail, Alert.

After resolving errors or making any modifications, it is necessary to re-run the 'Error Checking' command in the **ErrorChecker** worksheet (this is necessary to ensure that the Checker Report is updated to reflect the most recent changes in the template).

The template may not be accepted by the ISA if the Checker Report includes Fail error messages.

#### 14. SUBMITTING DATA

Data submissions are uploaded through the 'Upload' user interface on the ISA website: https://data.isa.org.jm/isa/map/

Large data submissions (greater than 20 MB) should not be submitted over the website user interface; contact the ISA Data Management Team to make other arrangements.

Subsequent to upload, the file status can be viewed in the Upload user interface on the website (e.g., 'Uploaded Successfully' indicates Template has been received, 'Archived' indicates the data has been added to the database or file library.

Submitted files must not have the same file name as previously submitted files.

##### 14.1 Resubmittals

When data from a previous submission are revised (i.e., errors were found and corrected, or the dataset was expanded) the data must be re-submitted. **Re-submissions must include the entire dataset, not just the portion of the data that were changed/revised.**

Also, data may have been submitted but not processed and included in DeepData (this can occur if the ISA Data Management Team find errors or issues that require the Contractor to update the file).

Re-submissions must not have the same name as previously submitted files. The file name should include a suffix indicating it is a revision. For example:

- original file name: BGRPMN12015_Geo_Template_2014rock.zip
- revised file name: BGRPMN12015_Geo_Template_2014rock_20210410rev.zip

The re-submitted file must be flagged as such in the Metadata Template by including a comment in the Remarks field.

Note that previous submissions will be removed from DeepData and archived; the re-submitted dataset will replace the original version. The archived data is not available over the DeepData website, but can be requested from the ISA.

#### 15. REFERENCES

International Seabed Authority (ISA). 2015. Consolidated regulations and recommendations on prospecting and exploration. Revised Edition.

International Seabed Authority (ISA). 2020. Recommendations for the guidance of contractors for the assessment of the possible environmental impacts arising from exploration for marine minerals in the Area. Issued by the Legal and Technical Commission, ISBA/25/LTC/6/Rev.1

Life Science Identifier (LSID) resolution project. (n.d.). Retrieved from http://www.lsid.info/

Horton T, Marsh L, Bett BJ, Gates AR, Jones DOB, Benoist NMA, Pfeifer S, Simon-Lledó E, Durden JM, Vandepitte L and Appeltans W (2021) Recommendations for the Standardisation of Open Taxonomic Nomenclature for Image-Based Identifications. *Front. Mar. Sci.* 8:620702. doi: 10.3389/fmars.2021.620702

Rabone M, Harden-Davies H, Collins JE, Zajderman S, Appeltans W, Droege G, Brandt A, Pardo-Lopez L, Dahlgren TG, Glover AG and Horton T (2019) Access to Marine Genetic Resources (MGR): Raising Awareness of Best-Practice Through a New Agreement for Biodiversity Beyond National Jurisdiction (BBNJ). *Front. Mar. Sci.* 6:520. doi: 10.3389/fmars.2019.00

#### Annex A

##### Listing of current ContractIDs to use when submitting data to DeepData.

| ContractID | Description |
|---|---|
| BGRPMN1 | Federal Institute for Geosciences and Natural Resources of Germany - PMN |
| BGRPMS1 | Federal Institute for Geosciences and Natural Resources of Germany - PMS |
| BPHDCPMN1 | Beijing Pioneer Hi-Tech Development Corporation - PMN |
| BRAZILCRFC1 | Companhia de Pesquisa de Recursos Minerais S.A. - CRFC |
| CIICPMN1 | Cook Islands Investment Corporation - PMN |
| CMMPMN1 | China Minmetals Corporation - PMN |
| COMRACRFC1 | China Ocean Mineral Resources Research and Development Association - CRFC |
| COMRAPMN1 | China Ocean Mineral Resources Research and Development Association - PMN |
| COMRAPMS1 | China Ocean Mineral Resources Research and Development Association - PMS |
| DORDPMN1 | Deep Ocean Resources Development Co. Ltd. - PMN |
| GSRPMN1 | Global Sea Mineral Resources NV - PMN |
| IFREMERPMN1 | Institut français de recherche pour l'exploitation de la mer - PMN |
| IFREMERPMS1 | Institut français de recherche pour l'exploitation de la mer - PMS |
| INDIAPMN1 | Government of India - PMN |
| INDIAPMS1 | Government of India - PMS |
| IOMPMN1 | Interoceanmetal Joint Organization - PMN |
| JOGMECCRFC1 | Japan Oil, Gas and Metals National Corporation - CRFC |
| KOREACRFC1 | Government of the Republic of Korea - CRFC |
| KOREAPMN1 | Government of the Republic of Korea - PMN |
| KOREAPMS1 | Government of the Republic of Korea - PMS |
| MARAWAPMN1 | Marawa Research and Exploration Ltd. - PMN |
| NORIPMN1 | Nauru Ocean Resources Inc. - PMN |
| OMSPMN1 | Ocean Mineral Singapore Pte. Ltd. - PMN |
| POLPMS1 | Government of the Republic of Poland - PMS |
| RUSFEDPMS1 | Government of the Russian Federation - PMS |
| RUSMNRCRFC1 | Ministry of Natural resources and environment of the russian federation - CRFC |
| TOMLPMN1 | Tonga Offshore Mining Limited - PMN |
| UKSRLPMN1 | UK Seabed Resources Ltd. - I - PMN |
| UKSRLPMN2 | UK Seabed Resources Ltd. - II - PMN |
| YUZHPMN1 | Yuzhmorgeologiya - PMN |

### Annex B

#### Example of coding unique Station IDs and Sample IDs for different types of sample collections.

| ContractID | Cruise Name | Leg Number | Station ID | Station Description | Sample ID | Sample Date | Actual Latitude | Actual Longitude | Upper Depth | Lower Depth | Depth Units | Sampling Device | Matrix Type | Vertical Datum | Remarks |
|---|---|---|---|---|---|---|---|---|---|---|---|---|---|---|---|
| **Example 1: Sampling a single sulphide chimney structure during 3 different ROV dives** | | | | | | | | | | | | | | | |
| IOMPMN1 | Yuzhmorgeologiya | 1 | ROV1 | Edifice 123 | Sample A | 4/21/2014 | 10.876133 | -119.450667 | 0 | 5 | cm | ROV | Sediment | SWI | sediment sampling |
| IOMPMN1 | Yuzhmorgeologiya | 1 | ROV2 | Edifice 123 | Sample A | 4/22/2014 | 10.865767 | -119.63215 | -9 | -9 | cm | ROV | Water filtered | NA | fluid sampling |
| IOMPMN1 | Yuzhmorgeologiya | 2 | ROV3 | Edifice 123 | Sample A | 9/2/2014 | 10.844667 | -119.83275 | 0 | 0 | cm | ROV | Rock | SWI | rock sampling |
| **Example 2: Drill sample** | | | | | | | | | | | | | | | |
| IOMPMN1 | Yuzhmorgeologiya | 2 | Core A | | Interval 1 | 5/15/2014 | 11.073117 | -119.661817 | 10 | 20 | cm | Drill | Sediment | SWI | |
| IOMPMN1 | Yuzhmorgeologiya | 2 | Core A | | Interval 2 | 5/15/2014 | 11.073117 | -119.661817 | 40 | 70 | cm | Drill | Sediment | SWI | |
| IOMPMN1 | Yuzhmorgeologiya | 2 | Core A | | Interval 3 | 5/15/2014 | 11.073117 | -119.661817 | 100 | 110 | cm | Drill | Sediment | SWI | |
| IOMPMN1 | Yuzhmorgeologiya | 2 | Core A | | Interval 4 | 5/15/2014 | 11.073117 | -119.661817 | 110 | 130 | cm | Drill | Sediment | SWI | |
| **Example 3: Different matrixes collected at the same Station** | | | | | | | | | | | | | | | |
| IOMPMN1 | Yuzhmorgeologiya | 2 | 1923 | | 01S | 5/24/2014 | 11.067583 | -119.65775 | 0 | 20 | cm | Gravity corer | Sediment | SWI | |
| IOMPMN1 | Yuzhmorgeologiya | 2 | 1923 | | 01PW | 5/24/2014 | 11.067583 | -119.65775 | 0 | 20 | cm | Gravity corer | Sediment Porewater filtered | SWI | |
| **Example 4: Total and dissolved fractions sampled from the same water sample.** | | | | | | | | | | | | | | | |
| IOMPMN1 | Yuzhmorgeologiya | 2 | 1942 | | 01T | 5/26/2014 | 11.0743 | -119.65455 | 3210 | 3211 | cm | Niskin bottle | Water unfiltered | AWI | |
| IOMPMN1 | Yuzhmorgeologiya | 2 | 1942 | | 01D | 5/26/2014 | 11.0743 | -119.65455 | 3210 | 3211 | cm | Niskin bottle | Water filtered | AWI | |
| **Example 5: Sediment collected from the same Station is sub-sampled and used for biological and chemisty testing** | | | | | | | | | | | | | | | |
| IOMPMN1 | Yuzhmorgeologiya | 1 | 3502 | | Sample A_bio | 5/30/2014 | 10.844667 | -119.83275 | 0 | 15 | cm | Box corer | Biological sample | SWI | |
| IOMPMN1 | Yuzhmorgeologiya | 1 | 3502 | | Sample A_chem | 5/30/2014 | 10.844667 | -119.83275 | 0 | 15 | cm | Box corer | Sediment | SWI | |

#### Annex C

**Example entries in the PhotoVideoFiles worksheet of the Metadata Template for various scenarios for collecting photos/videos.**

| | | | | | | |
|---|---|---|---|---|---|---|
| | 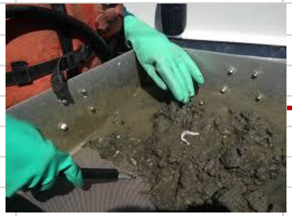 | 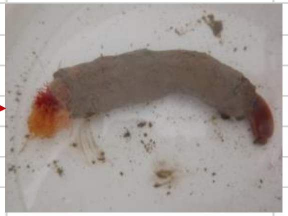 | 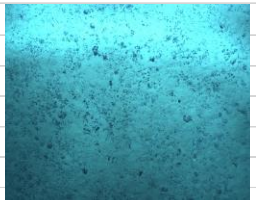 | 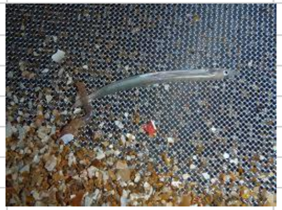 | 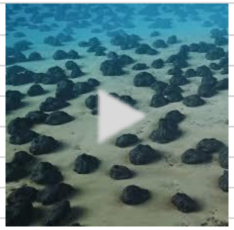 | 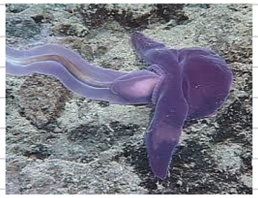 |
| **Scenario** | Scenario 1 | Scenario 2 | Scenario 3* | Scenario 4 | Scenario 5 | Scenario 6 |
| **Description of Scenario** | Photo of sample taken at a point location | Photo of organism taken from a point sample | Photo of the seafloor to record mineral abundance | Photos of organisms collected from trawl sample | Video track | Photo frames from video track |
| ***Field in PhotoVideoFiles worksheet*** | | | | | | |
| VP File Name | Prt001.jpg | ytt004.jpg | DSCN1005.jpg | DSCN2095.jpg | IMG_5854.mp4 | VidTrack1_211.mp4 |
| Short Title for Photo or Video File | Stn 1922-1 retrieved sample | Organism from Stn 1922-1 | seafloor nodules | Specimen A from 3546t trawl | Video for 3546t trawl | Organism B from 3546t traw |
| Associated Result | Sample - point | Individual - point | Sample - point | Individual - trawl | Sample - trawl | Individual - trawl |
| Station ID | 1922-1 | 1922-1 | ABC | -- | 3546t | 3546t |
| Sample ID | 1 | 1 | 1 | 57 | 57 | 57 |
| Trawl ID | -- | -- | -- | 3546t | -- | -- |
| OrgNum | -- | 3 | -- | -- | -- | 5 |
| Frame Code | -- | -- | -- | -- | -- | 456 |
| Area of Image | 0.2 | 0.2 | 0.5 | 0.2 | -- | -- |
| Height above seafloor | -- | -- | 4.2 | -- | 23 | 13 |
| Camera Specs | Nikon F mount 1.5 x lens focal width | Nikon F mount 1.5 x lens focal width | Nikon F mount 1.5 x lens focal width | Marshall CV502-WPMB Full HD 3.7 mm lens | Marshall CV502-WPMB Full HD 3.7 mm lens | Marshall CV502-WPMB Full HD 3.7 mm lens |
| Video/Photo Date/Time Stamp | 11/22/2014 14:15 | 11/22/2014 15:15 | 2/15/2014 17:05 | 11/27/2014 15:15 | 6/11/2014 15:20 | 6/11/2014 15:20 |
| Confidential Status | Public | Public | Confidential | Public | Public | Public |
| Confidentiality Duration | -- | -- | 5 | -- | -- | -- |
| Remarks | | | | | Date/time is the start time of video track | Date/time is the start time of video track |

*Photos that are not associated with samples or results may be submitted [e.g., a geological feature (black smoker), the seafloor, an organism swimming in the water column].
