## Supplementary material for "A review of the International Seabed Authority database DeepData: challenges and opportunities in the UN Ocean Decade": SF5A_Deep Data Darwin Core Mapping 2.pdf

20-Nov-2020

### Deep Data Correlation to Darwin Core in the Context of OBIS

This document outlines the correlation between Deep Data and Darwin Core for OBIS Integration. What is Darwin Core? According to https://dwc.tdwg.org/terms/ **Darwin Core is a body of standards for biodiversity informatics. It provides stable terms and vocabularies for sharing biodiversity data. Darwin Core is maintained by TDWG (Biodiversity Information Standards, formerly The International Working Group on Taxonomic Databases).**

### Deep Data

From deep data a number of structures were selected based on the definition of DwC Terms. These represent formatted input data submitted by Contractors for data on Biological samples collected at specific points or Biological Samples collected on a trawl

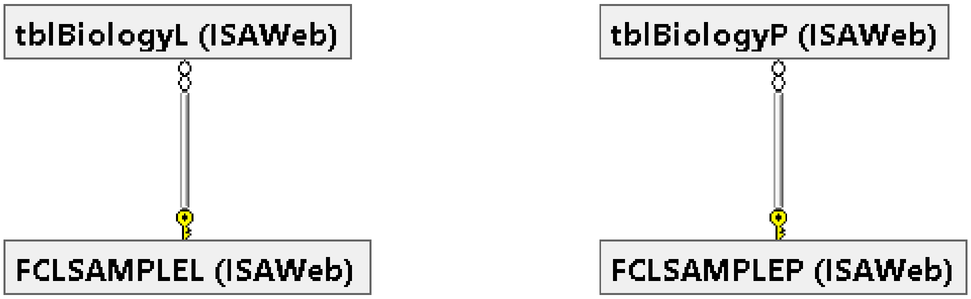


### DeepData Mapping Fields

Key mapping fields are taken from the ANALYSIS data.

| Category | Analysis |
|---|---|
| Ecology | Nominal size Category |
| Ecology | Number of individuals |
| Ecology | Relative abundance (%) |
| Ecology | Relative dominance (%) |
| Ecology | Species density - benthos (ind/m2) |

| | |
|---|---|
| Ecology | Species density - pelagic (ind/m3) |
| Ecology | Total Biomass collected (g/m2) |
| Organism Details | Sex |
| Taxonomist information | Taxonomist |
| Taxonomist information | Taxonomist E-mail |
| Taxonomist information | Taxonomist Institution |
| Taxonomy ID | Species |
| Taxonomy ID | Order |
| Taxonomy ID | Phylum |
| Taxonomy ID | Class |
| Taxonomy ID | Family |
| Taxonomy ID | Genus |
| Taxonomy ID | Identification Method |
| Taxonomy ID | Kingdom |
| Taxonomy ID | Morphotype |
| Taxonomy information | Database Taxa ID |
| Taxonomy information | Description of DNA Sequence |
| Taxonomy information | Notes on taxanomic identification |
| Taxonomy information | Putative species name or number |
| Taxonomy information | Voucher code |
| Taxonomy information | Taxonomic Database |
| Taxonomy information | Taxonomic Status |

### Darwin Core terms

DwC terms correspond to the column names of the dataset. A list of all possible Darwin Core terms can be found on TDWG. Below is an overview of the most relevant Darwin Core terms to consider when contributing to OBIS, with guidelines regarding their use.

Note that OBIS currently has eight required DwC terms: `occurrenceID`, `eventDate`, `decimalLongitude`, `decimalLatitude`, `scientificName`, `scientificNameID`, `occurrenceStatus`, `basisOfRecord`.

### Correlation Guidelines

ISA DeepData will not map all DwC Terms but will aim to provide as much comprehensive mapping as is available. Data currently unavailable with no relative mapping will be removed from final record dump provided to OBIS

#### Occurrence

| DwC Term | Deep Data Field |
|---|---|
| occurrenceID | Concatenation of TrawlID/StationID & SampleID |
| catelogNumber | Concatenation of Catalogue_OID & SampleID |
| RecordedBy | Concatenation of Taxonomist +','+Taxonomist E-mail |
| individualCount | Number of Individuals |

| | |
|---|---|
| organismQuantity | On condition if Species density has a value else Number of Individuals |
| organismQuantityType | In case of Species density then value: ‘indivdual density per metre cube’ else ‘individuals’ |
| sex | sex |
| occurenceStatus | ‘present’ |
| associatedSequences | Description of DNA Sequence |
| occurrenceRemarks | Remarks |

### Event

| DwC Term | Deep Data Field |
|---|---|
| eventID | Concatenation of Cruise leg and sampleID |
| eventDate | SampleDateStart/SampleDate |
| eventTime | Time part of SampleDate |
| month | Month part of SampleDate |
| day | Day part of SampleDate |
| Habitat | HabitatDescription |
| samplingProtocol | SamplingDevice |

### Location

| DwC Term | Deep Data Field |
|---|---|
| locationId | Concatenation of Station/TrawlID and AreaKey |
| MinimumDepthInMeters | minDepth |
| MaximumDepthInMeters | maxDepth |
| vermatimDepth | MinDepth and MaxDepth |
| decimalLatitude | StartLatidude |
| decimalLongitude | StartLongitude |
| verbatimCoordinateSystem | ‘decimal degrees’ |
| verbatimSRS | ‘WGS84’ |

### Identification

| DwC Term | Deep Data Field |
|---|---|
| identificationID | Concatenation of CruiseLeg and TaxonID |
| typeStatus | Taxonomic Status |
| identifiedBy | Taxonomist |
| SampleDateStart | dateIdentified |
| identificationVerificationStatus | 0 |
| typeStatus | Voucher status |

### Taxon

| DwC Term | Deep Data Field |
|---|---|
| taxonID | Concatenation of Cruise leg and SampleID |
| scientificName | Species or Genus |
| scientificNameID | Putative species name or number |
| Kingdom | Kingdom |
| Phylum | Phylum |
| class | Class |
| order | Order |
| family | Family |
| genus | Genus |
| taxonomicStatus | Taxonomic Status |

### RecordLevel

| DwC Term | Deep Data Field |
|---|---|
| type | 'Event' |
| license | Attribution 3.0 IGO (CC BY 3.0 IGO) |
| rightsHolder | Contractor |
| accessRights | Terms of use DeepData |
| bibliographicCitation | 'International Seabed Authority, DeepData' |
| institutionID | Interiational Seabed Authority |
| basisOfRecord | 'Taxon' |

In addition to the above mapping, data such as relative abundance is added to the MeasurementOrFact DwC standard. This can be extended to any variable in DeepData that can be represented.

### MeasurementOrFact

| DwC Term |
|---|
| measurementID |
| measurementType |
| measurementValue |
| measurementUnit |

### Data Cleaning

A number of steps were taken to cleanse the data provided by contractors in order to improve Taxon Matching on OBIS. This involved removing temporary names e.g. sp. and remap to

### File Transfer Protocol.

In providing data to OBIS, DeepData will be sending a single file per month of the complete dataset. Which will be picked up by OBIS Technical team and formatted for presentation on OBIS Platform.

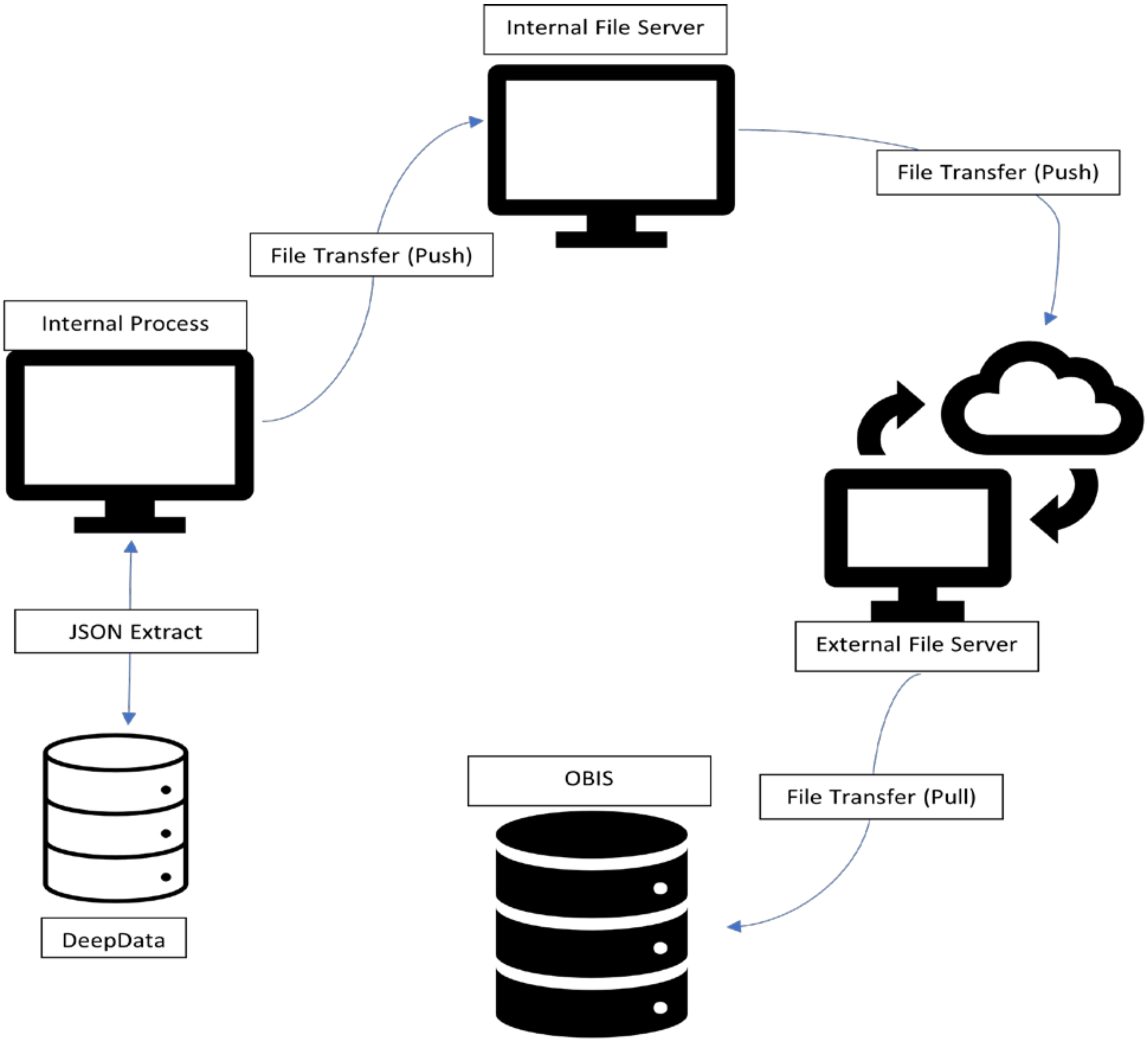
