## Supplementary material for "A review of the International Seabed Authority database DeepData: challenges and opportunities in the UN Ocean Decade": SF6_DeepData_review_data_processing_script.R

```
###########################################################
########### CCZ AND DEEPDATA ANALYSIS
setwd("C:\\Users\\murr\\Desktop\\EVERYTHING\\CURRENT_MANUSCRIPTS\\CCZ_and_DEEPDATA\\Data_analysis_CCZ_DeepData\\")

######## MATCH FILE NAMES

###################################

## load packages
library(dplyr)
library(tidyr)
library(robis)
library(curl) 
library(obistools)
library(devtools)
library(lubridate)
library(glmmTMB)
library(ggthemes)
library(DataExplorer)
library(gridExtra)
library(lme4)
library(car)
library(reshape2)
library(ggfortify)
library(boot)
library(aod)
library(mgcv)
library(pscl)
library(ggplot2)
library(RCurl)
library(stringr)
library(forcats)
library(readr)
library(glue)
library(xml2)
library(jsonlite)
library(purrr)
library(knitr)
library(uuid)
library(stringr)
library(worms)
library(viridis)
library(RColorBrewer)
library(GADMTools)
library(sp)
library(spatialEco)
library(spData)
library(maptools)
library(rgdal)
library(rgeos)
library(sf)
library(mapr)
library(leaflet)
library(rgbif)
library(rfishbase)
library(taxize)
library(scrubr)
library(mapr)
library(mregions)

###############################################################

## OBIS data collection
library(robis)

data.ccz <- occurrence(geometry = "POLYGON ((-160.3477 12.5906, -154.0898 -5.4997, 
                       -107.5430 8.6544, -121.8867 23.3343, -160.3477 12.5906))")

data.newer <-
  data.ccz %>%
  filter(depth >= 3000)

write.csv(data.newer, "OBIS_CCZ_3000m+_at_2021-07-12.csv")

##################################################

## SELECT COLUMNS

data1 <- read.csv("DD_PUBLISHED_4analysis_ed_2022-05-24.csv") 
data2 <- read.csv("DD_published_main_file_2021-07-12_v10.csv") 
merga <- merge(data1, data2, by="rec", all = T)
dim(merga) ## 511   7
dim(data2) ## 268   6
dim(data1) ## 511   2
write.csv(merga, "OUTPUT.csv")

DD <- read.csv("DD_published_main_file_2021-07-12_v10.csv")
names(DD)

DD_ed <- read.csv("DD_published_main_file_2021-07-12_v10.csv") %>% 
  dplyr::select(rec, comp_key, comp_key_temp, occurrenceID, occurrenceID_ed, Match.type, 
                Taxon.status, ScientificName_accepted, Phylum_ed,	Class_ed,	Order_ed,	Family_ed,	Genus_ed,
                Subgenus, Species_ed,	put_sp_ed,	put_cat,	put_valid,	scientificName,	taxonRank,
                TR_w_TCID,	qualifier,	rank_qual,	column_notes,	non_specimen,	incl_in_taxon_analysis,
                incl_strict_taxon_records, INCL,	CL_short_incl, sample_time,	SampleDate_ed,	sample_month,	
                sample_year, SampleCollectionMethod_ed, WaterDepth_ed, StorageLocation_ed, 
                HabitatType_ed, HabitatDescription_ed, ResearchVessel_ed,	Nominal.size.Category_ed, 
                Number.of.individuals_ed, ContractorID,	SubArea,	StationID, StationDesc,	SampleID,
                ActualLatitude,	ActualLongitude, AreaSampled, Phylum, Class, Order, Family, Subfamily,
                Species, Putative.species.name.or.number, Notes.on.taxanomic.identification,
                Taxonomic.Status, Description.of.DNA.Sequence,
                Voucher.code, Voucher.institution.code, Voucher.status,
                Relative.abundance...., Relative.abundance....) %>%
  dplyr::filter(INCL != "NO") %>% 
  mutate(SampleDate_ed = lubridate::dmy(SampleDate_ed), 
         #         WaterDepth = replace(WaterDepth, WaterDepth == "-9", "NA"),
         decimalLongitude = as.numeric(ActualLongitude),
         decimalLatitude = as.numeric(ActualLatitude)
  )

names(DD_ed)
dim(DD_ed) # 40521    65
head(DD_ed)

##################################################

## FINAL DATASET FOR ANALYSIS- remove rows to exclude, select relevant fields

obis.all <- read.csv("OBIS_CCZ_all_depths_2021-07-12_v12.csv")
dim(obis.all) ##    126329    216
names(obis.all)
obis <- obis.all%>% 
  dplyr::select(rec, layer, inclusion_criteria, inc_notes_mar2022, include, taxon_note, scientificName_ed,
                DD_record, date_year, year, ScientificName_accepted, qualif_ed, 
                TR_w_TCID, CCZ_endemic, other_rank, taxonRank_ed,
                Authority, AphiaID_accepted, occurrenceID, occurrenceID_ed,
                scientificName, decimalLatitude, originalScientificName, basisOfRecord, 
                minimumDepthInMeters, maximumDepthInMeters, depth, id, 
                class, order, family, decimalLongitude, decimalLatitude, occurrenceID, kingdom, phylum, rightsHolder,
                institutionID, individualCount, type, catalogNumber, samplingProtocol, eventID,
                taxonRank, eventTime, taxonomicStatus, verbatimDepth, locationID, accessRights, 
                habitat,recordedBy, identifiedBy, taxonRemarks, taxonomicStatus, genus, species, 
                subclass, taxonConceptID, identificationQualifier, previousIdentifications,
                institutionCode, recordNumber, datasetName, associatedSequences)%>% 
  dplyr::filter(include != "no") 
names(obis)
dim(obis) ## 50851    56

#####################################################

## select DeepData records in OBIS dataset

obis_DD <- obis %>% 
  filter(DD_record =="DD ISA records")
dim(obis_DD) ## 48554    56
write.csv(obis_DD, "OBIS_DD_4_analysis_2022-04-20.csv")

####################################################

## read in files

DD <- read.csv("DD_PUBLISHED_main_file_2021-07-12_v10.csv")
DD_ed <- read.csv("DD_PUBLISHED_4analysis_ed_2022-05-24.csv")
obis_DD <- read.csv("OBIS_DD_4_analysis_2022-04-20.csv")

dim(DD_ed)           # 40518  
dim(obis_DD)         # 48554  

##################################################

## MAPPING -OBIS- GBIF spatial join
## IN QGIS- create shpfile join of all the contract areas, reserved areas and APEIs
## convert to RDS file? not necessary- do readOGR
## could also do join directly in QGIS

zones <- readOGR(dsn = ".", layer = "CCZ_ISA_only_merged_file")
points <- read.csv("OBIS_CCZ_records_coords_2021-04-17.csv")
coordinates(points) <- ~longitude + latitude ### converted into SPDF, renamed decimalLatitude to latitude
class(points) ## SpatialPointsDataFrame # sp
head(points@coords)
head(points) 
points@bbox ## gives coords of bounding box
head(zones)
names(zones)
proj4string(points)
proj4string(zones) 
proj4string(points) <- CRS("+proj=longlat +datum=WGS84 +ellps=WGS84 +towgs84=0,0,0")
#Warning message:#In showSRID(uprojargs, format = "PROJ", multiline = "NO") :
#  Discarded datum WGS_1984 in CRS definition, but +towgs84= values preserved
points <- spTransform(points, CRSobj = CRS(proj4string(zones)))
#Warning message: In proj4string(zones) : CRS object has comment, which is lost in output
points_with_zones1 <- over(points, zones[,"layer"]) 
## produce a data frame with the same number of rows as your layer of points. 
## connect those two:

points$zonelayer <- points_with_zones1$layer
## replace "name" with the name of column containing zone names. 
## can also use column index
head(points_with_zones1)
write.csv(points_with_zones1, "GBIF_CCZ_SHP_MERGE_2021-04-17_v2.csv")
## worked

points <- read.csv("GBIF_CCZ_SHP_MERGE_2021-04-17_v2.csv")
## dataset with if in or out of TT, composite key, rec, lat long
points <- read.csv("OBIS_CCZ_SHP_MERGE_2021-04-17.csv") ## ie out CCZ
head(zones@data)  # polygons
head(points@data) # points
#Error in h(simpleError(msg, call)) : 
#error in evaluating the argument 'x' in selecting a method for function 'head': trying to get slot "data" from an object (class "data.frame") that is not an S4 object 
plot(zones)
points(points, pch=20)

##############################################

## Fig 2: taxonomic resolution of records by data source

data <- DD_ed%>%
  filter(other_rank != "y")

data <- obis_DD %>%
  filter(other_rank != "y")


p <- ggplot(data = data) + 
  geom_bar(mapping = aes(x = TR_w_TCID, fill =TR_w_TCID)) +
  labs(x = "Taxonomic rank", y = "Records")+ 
  #  labs(title="Taxonomic resolution of records: CCZ literature") +
  scale_fill_viridis_d() +
  theme(text=element_text(size=12), #change font size of all text
        axis.text=element_text(size=12), #change font size of axis text
        axis.title=element_text(size=12), #change font size of axis titles
        plot.title=element_text(size=12), #change font size of plot title
        legend.text=element_text(size=12), #change font size of legend text
        legend.title=element_text(size=12)) 
p_ed <- p + theme(legend.position = "none")
```
